# Antiglycation effects of imidazole dipeptides and 2-oxo-imidazole dipeptides on glyceraldehyde-induced intracellular protein glycation and neuronal cell death

**DOI:** 10.64898/2026.06.25.734660

**Authors:** Yasunari Yamada, Koji Hashida, Kohei Hayashi, Kenji Yoshimochi, Tsunehisa Hirose, Motoshi Shimotsuma, Yoshio Hamada, Kenji Usui, Naoki Yokoyama, Toshiaki Hara, Satoru Nishino, Hideaki Kakeya, Shozo Tomonaga, Makoto Ozaki

## Abstract

Glyceraldehyde (GA) contributes to the development of various diseases, such as diabetes and Alzheimer’s disease via protein glycation and the formation of advanced glycation end products (AGEs); however, effective strategies for neutralizing GA are limited. Carnosine (Car), an imidazole dipeptide (IDP) that is abundant in meat, suppresses protein glycation by scavenging reactive aldehydes. There are only a few reports on the antiglycation activity of Car against GA. For other IDPs, such as anserine, balenine (Bal), and homocarnosine, there are almost no reports on their antiglycation activity. In this study, we demonstrated the antiglycation activity of four types of IDPs and 2-oxocarnosine (2-oxo-Car), an oxidized form of Car, against GA-induced intracellular protein glycation and neuronal cytotoxicity. Car and Bal exhibited significantly higher reactivity with GA compared with other IDPs and 2-oxo-Car. An *in silico* analysis suggested that the difference in reactivity is dependent upon intramolecular hydrogen bond formation and the conformation of each IDP. Although there were differences in reactivity with GA, LC–MS analysis revealed that all of the IDPs and 2-oxo-Car reacted with two molecules of GA to form adducts containing pyridinium rings. Car and Bal exhibited high reactivity with GA and markedly suppressed GA-induced cytotoxicity in SH-SY5Y cells. Western blot and qPCR analyses revealed that IDPs suppressed GA-induced protein glycation and the upregulation of endoplasmic reticulum and oxidative stress response genes. Our results indicate that IDPs represent a novel preventive approach to AGE-related diseases and provide a foundation for the development of strategies to treat GA-related neurotoxicity.

**Graphical abstract:** Carnosine and balenine, which are imidazole dipeptides (IDPs), scavenged two molecules of glyceraldehyde to form adducts containing pyridinium rings and suppressed intracellular protein glycation and neuronal cell death. The formation of intramolecular hydrogen bond of IDPs played a crucial role in the strength of antiglycation activity.

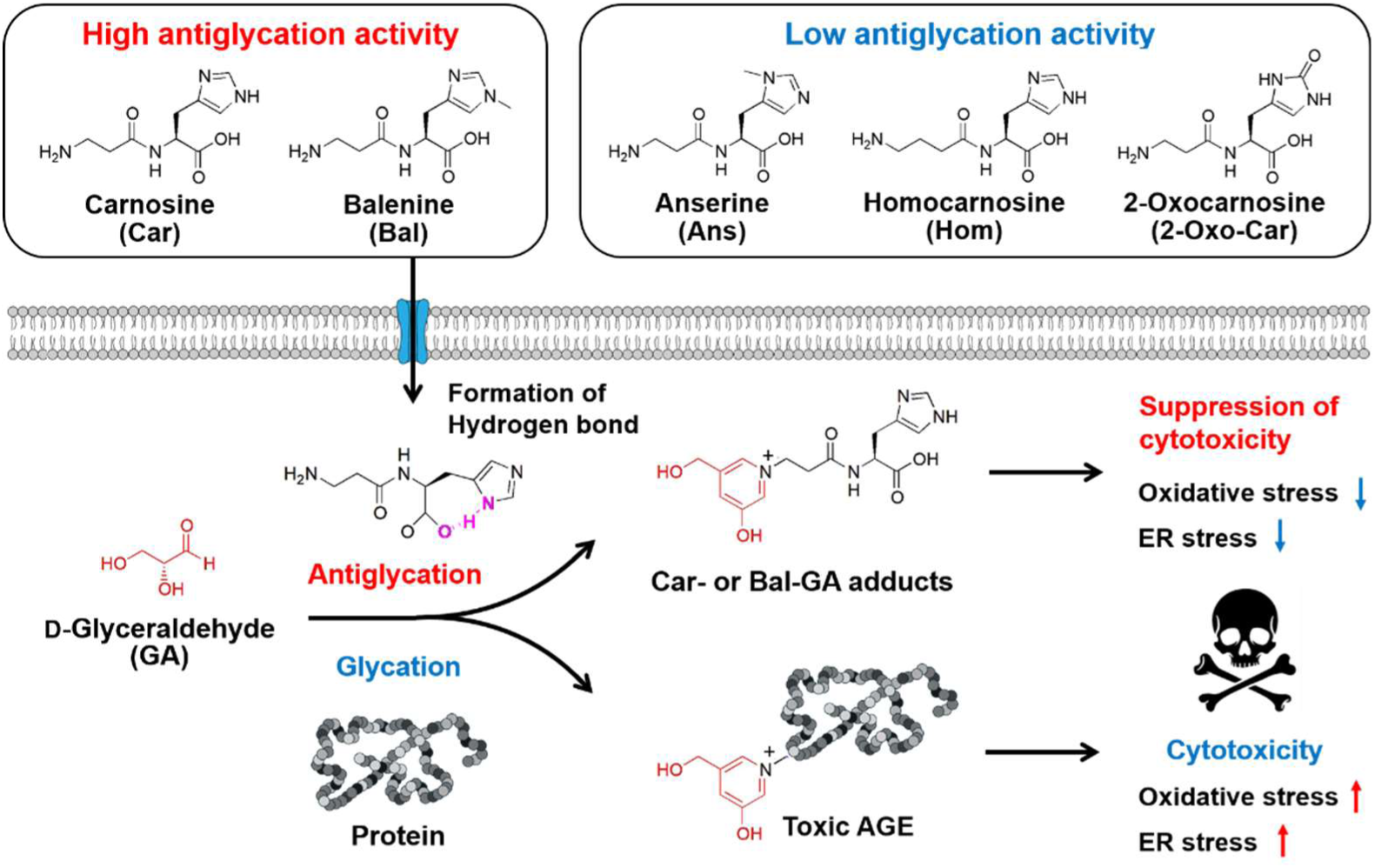

## Introduction

Glycation is a type of aging process that occurs when excess sugars ingested through the diet or reactive aldehydes produced via metabolism bind to proteins in the body to generate advanced glycation end products (AGEs), which are associated with aging. Various AGEs are produced depending on the reactive aldehydes that react with proteins. For example, *N^ε^*-(carboxymethyl)-lysine (CML) is produced by reacting with glyoxal and results from the oxidative cleavage of Amadori compounds and lipid peroxidation, with the amino group of the lysine (Lys) residue side chain.^1^ CML can also be produced by the reaction of glycolaldehyde, which is derived from hypochlorous acid and serine, with the amino group of the Lys residue side chain.^2^ In addition, the presence of reactive oxygen species (ROS), such as hydroxyl radical (OH·) and peroxynitrite (ONOO^-^), promotes CML formation.^3, 4^ Methylglyoxal hydroimidazolone 1 (MG-H1) is produced when methylglyoxal (MGO) reacts with the guanidino group of the arginine (Arg) residue side chain.^5^ MGO is produced from triose phosphate through a non-oxidative pathway in anaerobic glycolysis and modifies amino acids, nucleic acids, and proteins. In particular, AGEs derived from glyceraldehyde (GA) exhibit strong cytotoxicity and are known as toxic AGEs (TAGEs). TAGE, which has a pyridinium ring, is produced when the amino groups of the Lys side chain of the protein react with two molecules of GA (Fig. 1A).^6, 7^ The accumulation of TAGE is involved in various diseases, such as diabetes, metabolic dysfunction-associated steatohepatitis, cardiovascular disease, Alzheimer’s disease, and cancer.^8–11^ Therefore, minimizing the accumulation of TAGE produced in the body is important for the prevention of these serious diseases.

**Fig. 1.**
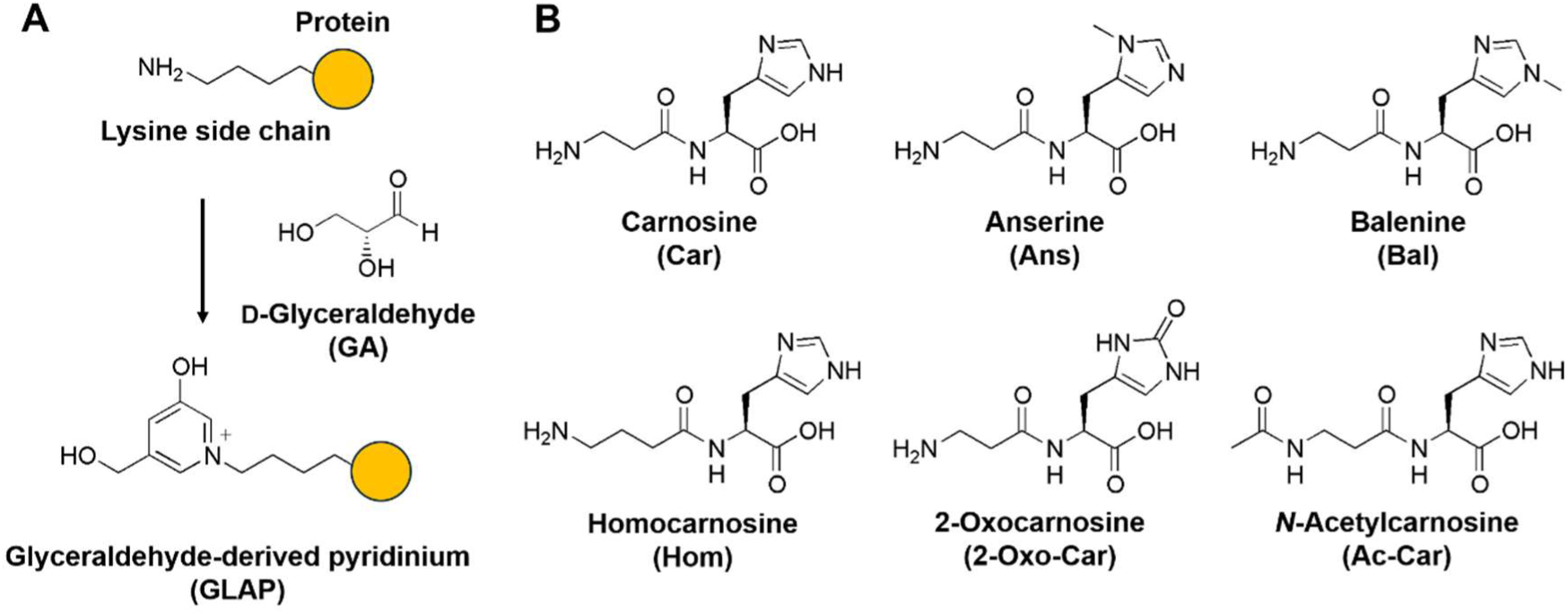
(A) Schematic illustration of GLAP generated from the amino group of the lysine residue side chain in protein and GA. (B) Chemical structures of Car, Ans, Bal, Hom, 2-oxo-Car, and Ac-Car used in this study.

Imidazole dipeptides (IDPs, Fig. 1B) such as carnosine (*β*-alanyl-L-histidine, Car), anserine (*β*-alanyl-*N^π^*-methyl-L-histidine, Ans), balenine (*β*-alanyl-*N^τ^*-methyl-L-histidine, Bal), and homocarnosine (*γ*-aminobutyryl-L-histidine, Hom) are functional food ingredients found in meat.^12–15^ They have attracted interest because they exert physiological activities that are beneficial to animals, such as antioxidant and anti-fatigue effects.^13, 16–18^ Recently, Ihara *et al*. demonstrated that 2-oxocarnosine (2-oxo-Car, Fig. 1B), in which the 2-position of the imidazole group is oxidized, occurs in trace amounts in meats and various tissues.^19–22^ 2-Oxo-Car exhibits significantly higher antioxidant activity compared with Car,^21, 23^ and markedly suppresses tyrosine nitration and intracellular ROS generation induced by rotenone and pyocyanin;^21, 24, 25^ however, there are few reports on the function of 2-oxo-Car other than its antioxidant activity. Car exhibits antiglycation effects resulting from its reactivity with various toxic aldehydes. Specifically, it captures *α*- and *β*-unsaturated aldehydes [4-hydroxy-2-nonenal (4-HNE), acrolein, and trans-2-hexanal] and dicarbonyl compounds [MGO and malondialdehyde (MDA)], which results in the formation of non-toxic and stable adducts;^26–29^ however, the reactivity and antiglycation effect of Car with GA, a precursor of TAGE that exhibits strong cytotoxicity, remains unclear. Because the reaction between Car and reactive aldehydes is a multi-step reaction involving the primary amine and the imidazole ring, other IDPs and 2-oxo-Car may exhibit different reactivity compared with Car; however, there are few reports on the aldehyde scavenging ability of IDPs other than Car. Ans, Bal, Hom, and 2-oxo-Car are more resistant to degradation by carnosinase (Car-degrading enzyme) compared with Car.^30, 31^ Therefore, it is important to evaluate their antiglycation activity because they may exhibit higher bioavailability than Car, thus representing a novel approach to preventing AGE-related diseases.

In this study, we evaluated the antiglycation effects of IDPs and 2-oxo-Car against GA *in vitro* and *in cellulo*. Specifically, the reactivity of each IDP and 2-oxo-Car with GA was measured using high-performance liquid chromatography (HPLC), and the structures of the reaction products and binding abilities were determined by liquid chromatography–mass spectrometry (LC–MS) and simulation analysis using Conformational Search. In addition, the antiglycation effects of IDPs and 2-oxo-Car in suppressing GA-induced intracellular protein glycation and cytotoxicity in SH-SY5Y cells were evaluated. Furthermore, the detailed mechanism of neuronal cell death induced by GA through quantitative polymerase chain reaction (qPCR) analysis was examined.

## Results and discussion

### Confirmation of IDP reagent purity

For the accurate determination of the antiglycation effect of IDPs and 2-oxo-Car, it is important to use high-purity compounds. Several studies have reported that IDPs obtained from several manufacturers are contaminated with other IDPs or 2-oxo-IDPs, which can affect their functionality.^23, 25^ We previously identified high-purity Car, Ans, Bal, and 2-oxo-Car reagents without other IDPs and 2-oxo-IDPs by DPPH radical scavenging assay, HPLC, and LC–MS measurements;^25^ however, the purity of the Hom reagent was not confirmed. Therefore, the purity of the Hom reagent was evaluated by HPLC and LC–MS. The peak positions of Hom, other IDPs, and 2-oxo-Hom were identified using our previously described methods (Fig. S1A).^15, 25^ The peaks derived from other IDPs and 2-oxo-Hom were not observed in the Hom reagent used in the present study, either through HPLC or LC–MS (Fig. S2A, S2B, and S3). These results indicate that IDPs and 2-oxo-Car reagents do not contain any other IDPs or 2-oxo-IDPs.

### Reactivity of IDPs and 2-oxo-Car with GA

The reactivity of each IDP and 2-oxo-Car with GA was evaluated using HPLC. For all samples, except *N*-acetyl-Car (Ac-Car, Fig. 1B), in which the amino group of Car is acetylated, the peaks derived from each IDP and 2-oxo-Car were reduced compared with the samples without GA, and new peaks were detected between 6 and 11 min (Fig. 2A). For Ac-Car, no decrease in the peak area value was observed with or without the addition of GA (Fig. S4). These results suggest that GA does not react with imidazole groups but reacts with amino groups. The reaction rate was calculated from the decrease in the peak area of each IDP and 2-oxo-Car in the samples with GA relative to the control sample. Car and Bal (60.6% and 62.8%) exhibited significantly higher reaction rates with GA compared with other IDPs (32.2%–45.7%), respectively (Fig. 2B). The difference in peak area values detected between 6 and 11 minutes in each sample correlated with the reaction rate, suggesting that it was a reaction product with GA (Fig. 2C). Next, the binding mode of each IDP and 2-oxo-Car with GA was analysed and identified using LC–MS. The peaks detected between 6 and 11 min were *m*/*z* 335.1 (Car), *m*/*z* 349.1 (Ans, Bal, and Hom), and *m*/*z* 351.1 (2-oxo-Car) (Fig. S5). The difference in *m*/*z* values was consistent with the difference in molecular weight (Car: 226.2; Ans, Bal, and Hom: 240.3; 2-oxo-Car: 242.2) between each imidazole dipeptide and 2-oxo-Car, suggesting that these peaks belong to IDP-GA adducts. A voltage was applied to the samples in the Q-array mode attached to the LC–MS, and the structure of each reaction product was identified from the resulting fragment pattern. For each reaction product, the fragments observed in the IDPs or 2-oxo-Car alone (Car: *m*/*z* 110.1; Ans: *m*/*z* 109.1; Bal: *m*/*z* 109.1 and 124.1; Hom: *m*/*z* 110.1; 2-oxo-Car: *m*/*z* 109.1 and 126.1, Fig. S6 and S7) and other characteristic fragments (Car, Ans, Bal, and 2-oxo-Car: *m*/*z* 198.1; Hom: *m*/*z* 178.1) derived from IDP-GA adducts were detected (Fig. 2D, E). GA reacts with Lys and Arg residues in various proteins to form TAGEs, such as GA-related pyridinium (GLAP) and argpyrimidine.^32^ These AGEs also react with two molecules of GA to form a pyridinium ring structure. We found that the N-terminal amino groups of each IDP and 2-oxo-Car bind to two molecules of GA to form a pyridinium ring, similar to the AGEs (Fig. 2F).

**Fig. 2.**
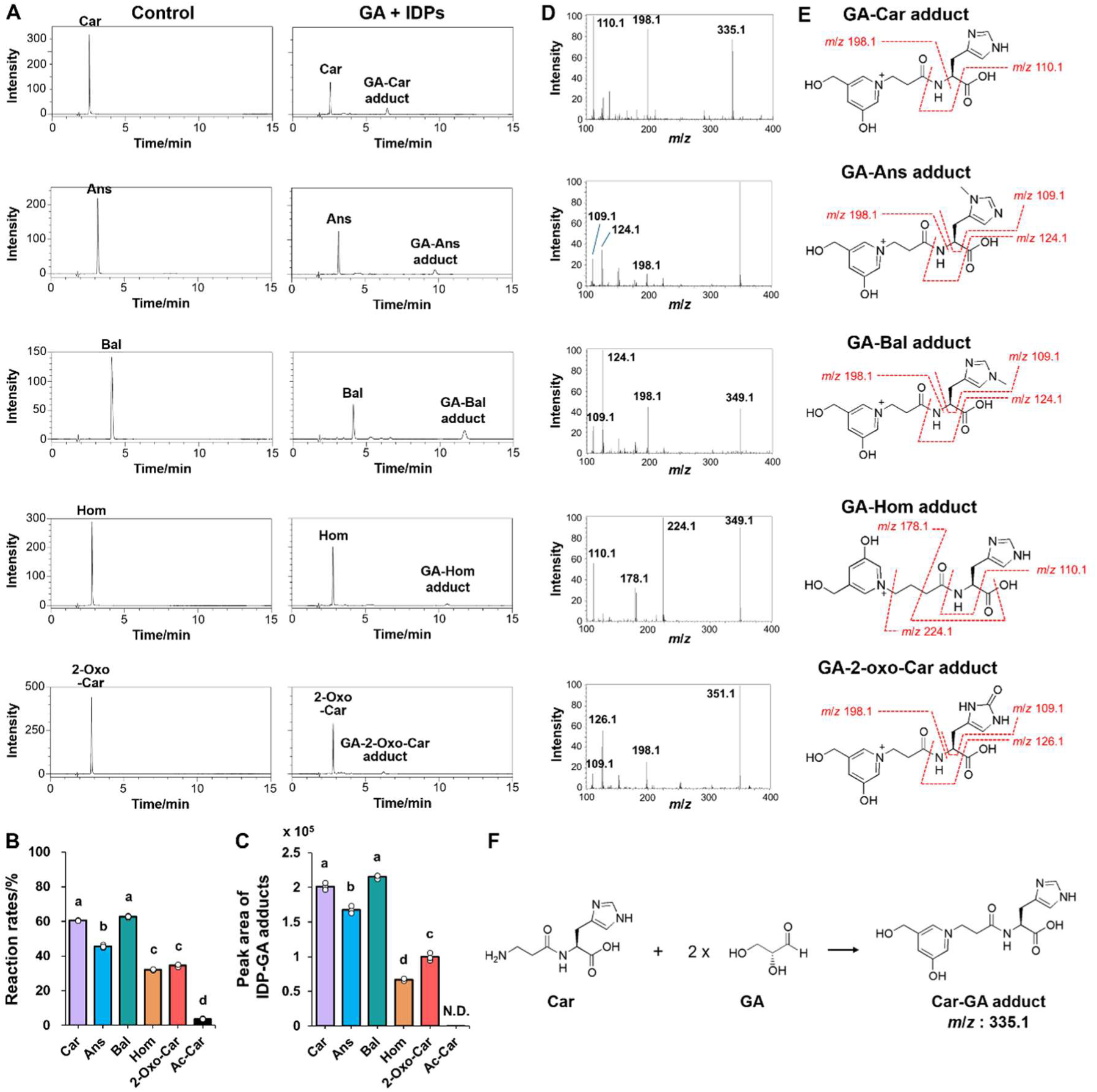
(A) HPLC chromatograms for the samples after incubating each IDP alone or 2-oxo-Car alone and each IDP or 2-oxo-Car with GA for 24 h at 37°C. (B) The reaction rates of each IDP and 2-oxo-Car with GA were calculated from the peak area values of the HPLC chromatograms.(C) The peak area values of each IDP-GA adduct obtained by HPLC. The data are presented as the mean ± SD (N = 3). Different letters indicate statistically significant differences among groups based on a one-way ANOVA followed by Tukey’s post hoc test (*p* < 0.05). (D) MS spectra of IDPs-GA and 2-oxo-Car-GA adducts obtained by LC–MS analysis. (D) Chemical structures of each IDP-GA adduct were identified based on MS spectra. (F) Reaction scheme of Car and two molecules of GA.

### Determination of the inhibitory effects of IDPs on AGEs formation

The glycation of BSA induced by GA and the inhibitory effects of IDPs and 2-oxo-Car against glycation were evaluated by sodium dodecyl sulfate-polyacrylamide gel electrophoresis (SDS-PAGE), followed by Coomassie brilliant blue (CBB) staining and western blot analysis using an anti-GA-pyridine antibody. No differences were observed among the samples by SDS-PAGE and CBB (Fig. S8A, B). In contrast, western blot analysis using an anti-GA pyridine antibody revealed that the band intensities of samples treated with Car and Bal were reduced compared with the control (GA alone) (Fig. 3A and S9). The AGE generation inhibition rates of Car and Bal were 72.3% and 68.2%, respectively (Fig. 3B). These results indicate that Car and Bal inhibit the formation of GLAP (e.g., BSA glycation). The samples treated with Ans (26.2%) slightly inhibited the formation of GLAP, whereas Hom and 2-oxo-Car (<5%) showed no inhibition (Fig. 3B). Interestingly, L-histidine (His) and *β*-alanine (*β*Ala), the constituent amino acids of Car, did not significantly inhibit GLAP formation (22.8%, Fig. 3B). The inhibitory effect of each IDP on the glycation of BSA depended on the reactivity of each IDP with GA. These results indicate that IDPs inhibit GA-induced protein glycation by directly reacting with GA. Furthermore, not all compounds containing an amino group can inhibit GLAP formation, as the specific conformation of imidazole dipeptides is important for inhibition.

**Fig. 3.**
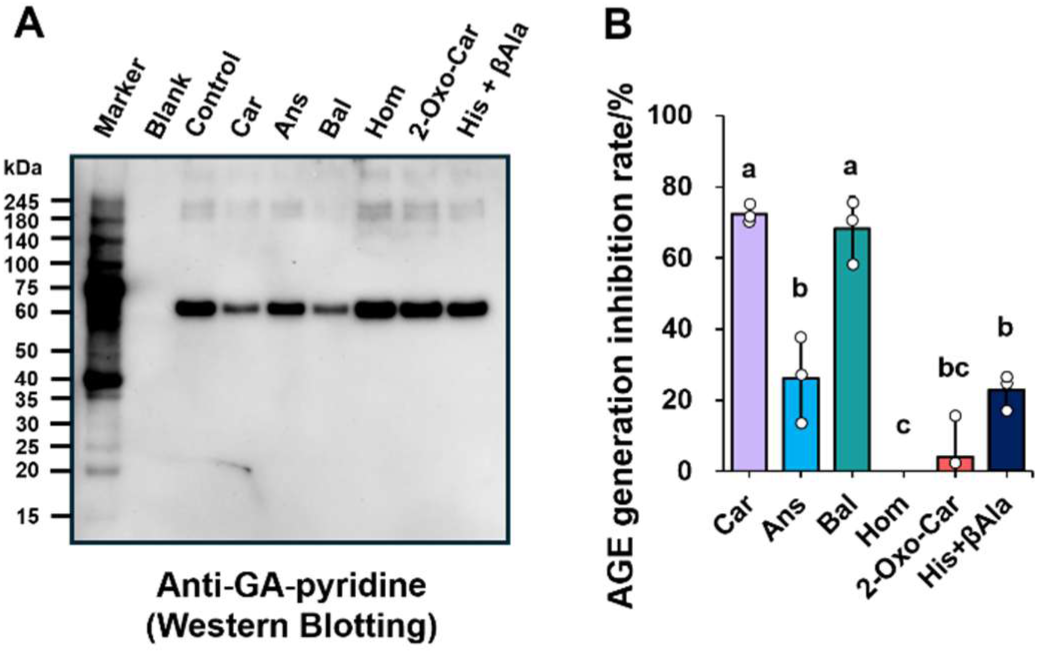
(A) Glycation level of BSA after incubating GA with each IDP, 2-oxo-Car, or His and *β*Ala for 24 h at 37°C. (B) Inhibition rate of AGE generation for each IDP, 2-oxo-Car, or His and *β*Ala, quantified based on western blot analysis. The data are presented as the mean ± SD (N = 3). Different letters indicate statistically significant differences among groups based on a one-way ANOVA followed by Tukey’s post hoc test (*p* < 0.05).

### Protective effects of IDPs against GA-induced cytotoxicity

In human neuroblastoma SH-SY5Y cells, GA induces intracellular TAGE formation and exerts neurotoxic effects.^33^ Thus, the protective effect of IDPs against GA-induced cytotoxicity was examined using cell viability and lactate dehydrogenase (LDH) assays. Microscopic examination of the morphology of SH-SY5Y cells treated with GA for 48 h revealed a decrease in the number of viable cells and an increase in dead cells; however, co-treatment with Car or Bal markedly restored the number of viable cells (Fig. 4A). In the cell count assay, cell viability was significantly reduced by GA (control, 55.4%), whereas Car (93.1%) and Bal (97.6%) significantly attenuated the decrease in cell viability resulting from GA treatment (Fig. 4B). Although Ans (80.3%) did not significantly affect cell viability, it slightly increased cell viability compared with the control (Fig. 4B). In contrast, no improvement in cell viability was observed in cells treated with 2-oxo-Car (57.8%), Hom (55.0%), or His + *β*Ala (54.2%). In the LDH assay, Car and Bal had an absorbance similar to that of the blank (Fig. 4C). Cells treated with Hom and 2-oxo-Car, which showed low cell viability, exhibited LDH-derived absorbance values similar to those of the control. These results indicate that Car and Bal attenuate GA-induced cytotoxicity.

**Fig. 4.**
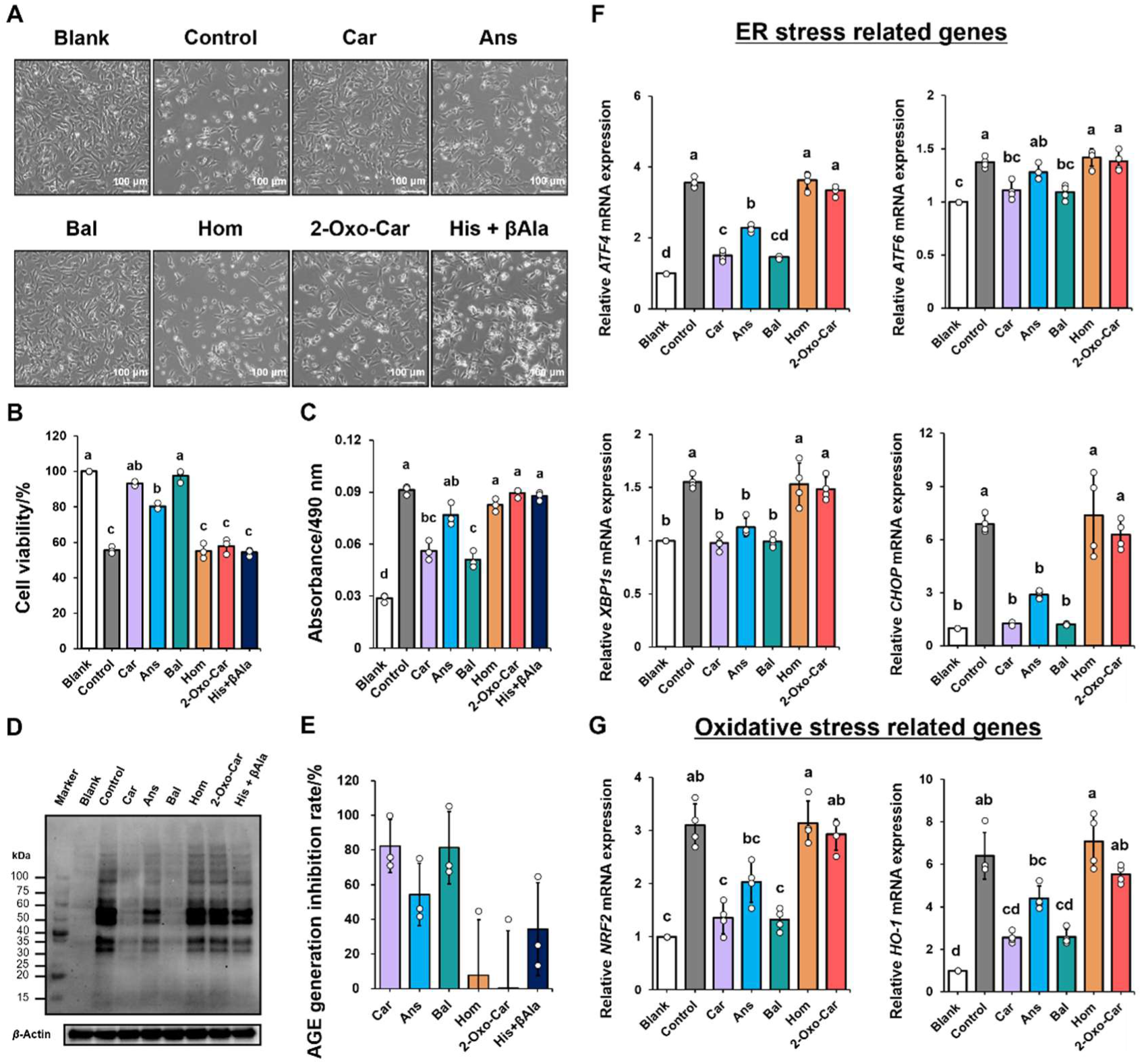
(A) Microscopic images and (B) cell viability of SH-SY5Y cells following treatment with GA and IDPs, 2-oxo-Car, or His and *β*Ala for 48 h at 37°C in a 5% CO_2_ atmosphere. (C) LDH activity in the culture medium of each sample was measured at 490 nm using an LDH assay kit. (C) Glycation level of intracellular proteins treated with GA and each IDP, 2-oxo-Car, or His and *β*Ala after incubating for 24 h at 37°C in a 5% CO_2_ atmosphere. (E) AGE generation inhibition rate of each IDP, 2-oxo-Car, or His and *β*Ala quantified by western blot analysis. The data are presented as the mean ± SD (N = 3). SH-SY5Y cells were treated with 800 μM GA and 2 mM IDPs or 2-oxo-Car for 24 h, followed by the measurement of (F) *ATF4*, *ATF6*, *XBP1s*, and *CHOP* ER stress-related gene expression and (G) *NRF2*, and *HO-1* oxidative stress-related gene expression by qPCR. The data are presented as the mean ± SD (N = 4). Different letters indicate statistically significant differences among groups based on a one-way ANOVA followed by Tukey’s post hoc test (*p* < 0.05).

Glycation of the intracellular proteins induced by GA and the inhibitory effect of IDPs and 2-oxo-Car against glycation were examined by SDS-PAGE analysis followed by western blot analysis using an anti-GA-pyridine antibody. The accumulation of intracellular GLAP was increased by GA treatment, but was markedly suppressed by Car and Bal (Fig. 4D and S10). Furthermore, when SH-SY5Y cells were treated with 0.25–2 mM Car, Car suppressed the formation of intracellular GLAP in a concentration-dependent manner (Fig. S11 and S12). Ans and His + *β*Ala slightly inhibited GLAP generation, while Hom and 2-oxo-Car did not inhibit (Fig. 4D and S10). Cell viability depends on the reactivity of GA with each IDP and the level of intracellular GLAP accumulation, which suggests that Car and Bal directly react with GA to suppress TAGE accumulation and protect cells.

Several studies have indicated that Car protects cells from the toxicity of aldehydes, such as 4-HNE, MDA, acetaldehyde, and MGO in various cell types;^26–29, 34^ however, the antiglycation activity of GA, which exhibits strong cytotoxicity in nerve cells, is unknown. The relationship between other IDPs, such as Ans, Bal, Hom, and 2-oxo-Car, with aldehyde-induced cytotoxicity has not been determined. The present study revealed that Car and Bal inhibit the accumulation of GA-derived AGEs, which are highly toxic, whereas other IDPs do not show antiglycation activity against GA. TAGE is localized to the neurons of the hippocampus and parahippocampal gyrus in AD patients.^35^ Moreover, TAGE induces the abnormal aggregation of *β*-tubulin and tau phosphorylation.^36, 37^ Therefore, intracellular TAGE may be a causative agent in the onset and progression of AD. Hom, which is abundant in the brain, may react with the carbonyl toxin acrolein and exhibit a detoxifying effect similar to that of Car.^38^ Interestingly, Hom did not react with GA and did not reduce its cytotoxicity. These results suggest that Car, not Hom alone, may be involved in neuroprotective and anti-inflammatory effects in the brain. Bal is more resistant to degradation by carnosinase compared with Car and has high bioavailability.^30, 31^ Therefore, it may be used as an alternative supplement to Car after clarifying the details of its metabolism and kinetics. Compounds that exhibit antiglycation effects against GA include L-carnitine and aminoguanidine. L-Carnitine has a protective effect against GA-induced neuronal injury by retaining mitochondrial function; however, the protective effect is only approximately 20%.^39^ In contrast, Car and Bal significantly inhibited GA-induced cell death (approximately 30–40%). This study suggests that IDPs may be superior to existing anti-aldehyde compounds in terms of neuroprotection against GA-induced nerve injury. Aminoguanidine, which was developed as a treatment for diabetic complications, also exerts a strong antiglycation effect; however, safety and efficacy issues are associated with long-term administration. Aminoguanidine is toxic to SH-SY5Y cells at 500 μM;^40^ however, IDPs exhibited no cytotoxicity, even at 2 mM. This suggests that IDPs may represent safe and effective antiglycation compounds.

### GA-induced stress response and protective effect of IDPs in SH-SY5Y cells

To elucidate the molecular mechanisms underlying the protective effect of IDPs against GA-induced toxicity, we measured the expression of ER stress-related [*ATF4*, *ATF6*, spliced *XBP1* (*XBP1s*), and *CHOP*] and oxidative stress-related (*NRF2* and *HO-1*) genes by qPCR. Cells treated with GA exhibited marked upregulation of *ATF4*, *ATF6*, *XBP1s*, *CHOP*, *NRF2*, and *HO-1*, whereas co-treatment with Car or Bal significantly suppressed their expression. GA is highly reactive and may disrupt protein homeostasis by modifying Lys and Arg residues in proteins, leading to ER stress. In the present study, the upregulation of *CHOP* and unfolded protein response (UPR)-related transcription factors (*ATF4*, *ATF6*, and *XBP1s*) was observed, suggesting that GA exposure activates all three pathways of the ER stress response (ATF6, IRE1, and PERK pathways).^[41]^

In addition to ER stress, GA exposure activated the oxidative stress response, as evidenced by *NRF2* and *HO-1* upregulation. *NRF2* activation likely reflects an adaptive antioxidant response to GA-induced redox imbalance. ER stress and oxidative stress pathways are closely related, and their reciprocal amplification may contribute to cytotoxicity. Interestingly, these stress responses were markedly attenuated by Car and Bal, suggesting that its protective effect is associated with suppression of GA-triggered cellular stress signaling. Car and Bal reacted with GA and suppressed the accumulation of intracellular proteins (Fig. 4D, E). These results suggest that Car and Bal act not by directly inhibiting downstream stress signaling pathways, but primarily reduce the availability of GA upstream. In contrast, *RAGE* expression decreased following GA exposure (Fig. S13). Although the cell types differed, several studies indicated that treating cells with AGEs increases the expression of *RAGE* itself through the activation of RAGE.^42–44^ In the present study, GA treatment did not upregulate the expression of *RAGE*. This may have occurred because GA, a precursor of AGEs, forms AGEs after cell uptake, thereby inducing cytotoxicity without involving RAGE on the cell surface. In addition, although there was a difference in *RAGE* expression between Car and Bal, its expression was independent of intracellular GLAP accumulation levels and cell viability. These results suggest that the cytotoxic effects observed in the present study are not likely mediated through activation of the canonical AGE-RAGE signaling pathway.

### *In silico* analysis of IDPs and 2-oxo-Car using Conformational Search

The differences in reactivity between each IDP and 2-oxo-Car with GA may be related to differences in their conformation in solution. Therefore, the stable conformations of each IDP and 2-oxo-Car in solution were determined by *in silico* analysis using Conformational Search. Car and Bal, which showed high reactivity with GA, contain a protonated nitrogen at the *N^π^* position of the imidazole rings and the C-terminal carboxyl group of the main chain in proximity to form a hydrogen bond (Fig. 5A). Under neutral conditions, Car is a tautomer in which the nitrogen at the *N^τ^* or *N^π^* position of the imidazole ring is protonated (Fig. 5B).^13, 45^ The distinct electronic and hydrogen-bonding capabilities of a tautomer result in significant differences in their effects on peptides. Li *et al*. reported that the nitrogen at the *N^π^* position of the imidazole group is protonated, causing this site to function as a hydrogen bond donor.^46^ For this reason, Car and Bal form a hydrogen bond between the protonated nitrogen at the *N^π^* position and the C-terminal carboxyl group adjacent to it. In Ans, the C-terminal carboxyl and imidazole groups did not form a hydrogen bond because of steric hindrance from the methyl group at the *N^π^* position of the imidazole ring (Fig. 5A). Hom is a molecule in which *β*Ala of Car is replaced by GABA; therefore, its main chain was longer than that of Car (Fig. 1B). In Hom, the N-terminal amino group formed hydrogen bonds at three points with the C-terminal carboxyl group and the protonated nitrogen at the *N^π^*position of the imidazole ring (Fig. 5A). Based on the calculated interatomic distances (Car: 3.26 Å; Hom: 2.27 Å, Fig. S14), hydrogen bonding between the N-terminal amino group and the C-terminal carboxyl group was observed only in Hom. Hydrogen bonding between the imidazole group of His and the carboxyl group of the Asp side chain in proteins affects the proton donor and acceptor properties. His-Asp pairs linked by a hydrogen bond function as proton relay systems during the catalysis of lactate dehydrogenase and malate dehydrogenase.^47^ Therefore, Car and Bal, which formed hydrogen bonds, exhibited higher reactivity with GA compared with that of other IDPs. Furthermore, in Bal, the hyperconjugated structure of the methyl group of methylated histidine is thought to contribute to the stabilization of the histidine cation. Some artificial enzymes have been reported that contain methylhistidine as a catalytic group in their active site.^48–52^ While the presence of a methyl group is thought to weaken the activity, this hyperconjugated structure may play some role in the strong activity of Bal. In Hom, the N-terminal amino group was used for hydrogen bonding, suggesting a decrease in GA reactivity. In 2-oxo-Car, the protonated nitrogen at the *N^π^* position and the C-terminal carboxyl group formed a hydrogen bond (Fig. 5A). The low reactivity with GA is likely due to the electron-withdrawing carbonyl oxygen at the 2-position of the imidazole ring. Furthermore, in Ac-Car, the protonated nitrogen at the *N^π^* position and the C-terminal carboxyl group formed a hydrogen bond (Fig. 5A); however, the N-terminal amino group, which is the reaction site with GA, was acetylated, resulting in a lack of reactivity with GA. These results suggest that hydrogen bond formation and the conformation of peptides influence the reactivity of each IDP and GA. In summary, Car and Bal, which form a hydrogen bond, scavenge two molecules of GA through a proton relay system, thereby suppressing GA-induced protein glycation and neuronal cell death (Fig. 6).

**Fig. 5.**
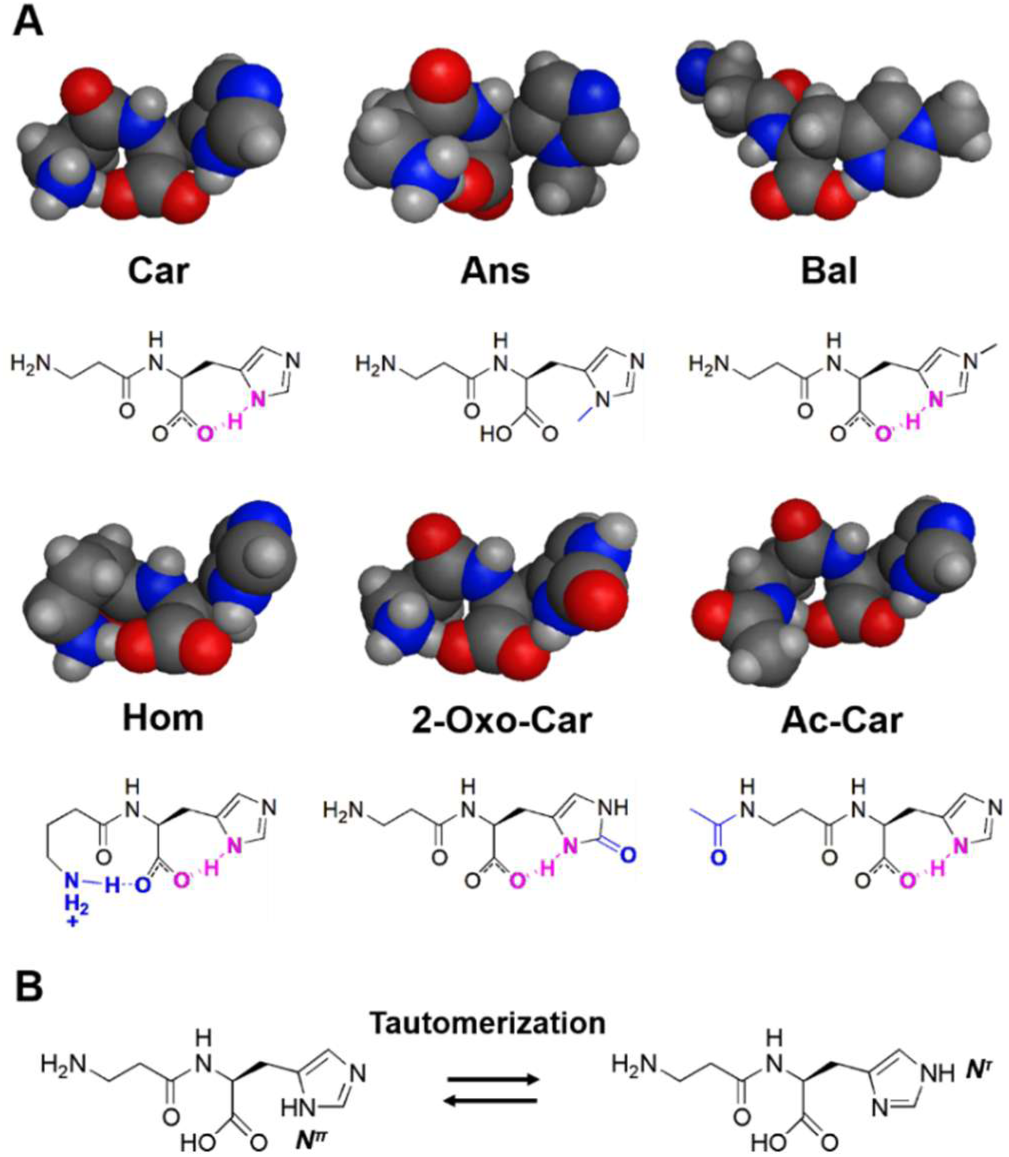
(A) Stable structures of Car, Ans, Bal, Hom, 2-oxo-Car, and Ac-Car in solution obtained using *in silico* analysis with Conformational Search. (B) Tautomeric forms of the imidazole ring of Car.

**Fig. 6.**
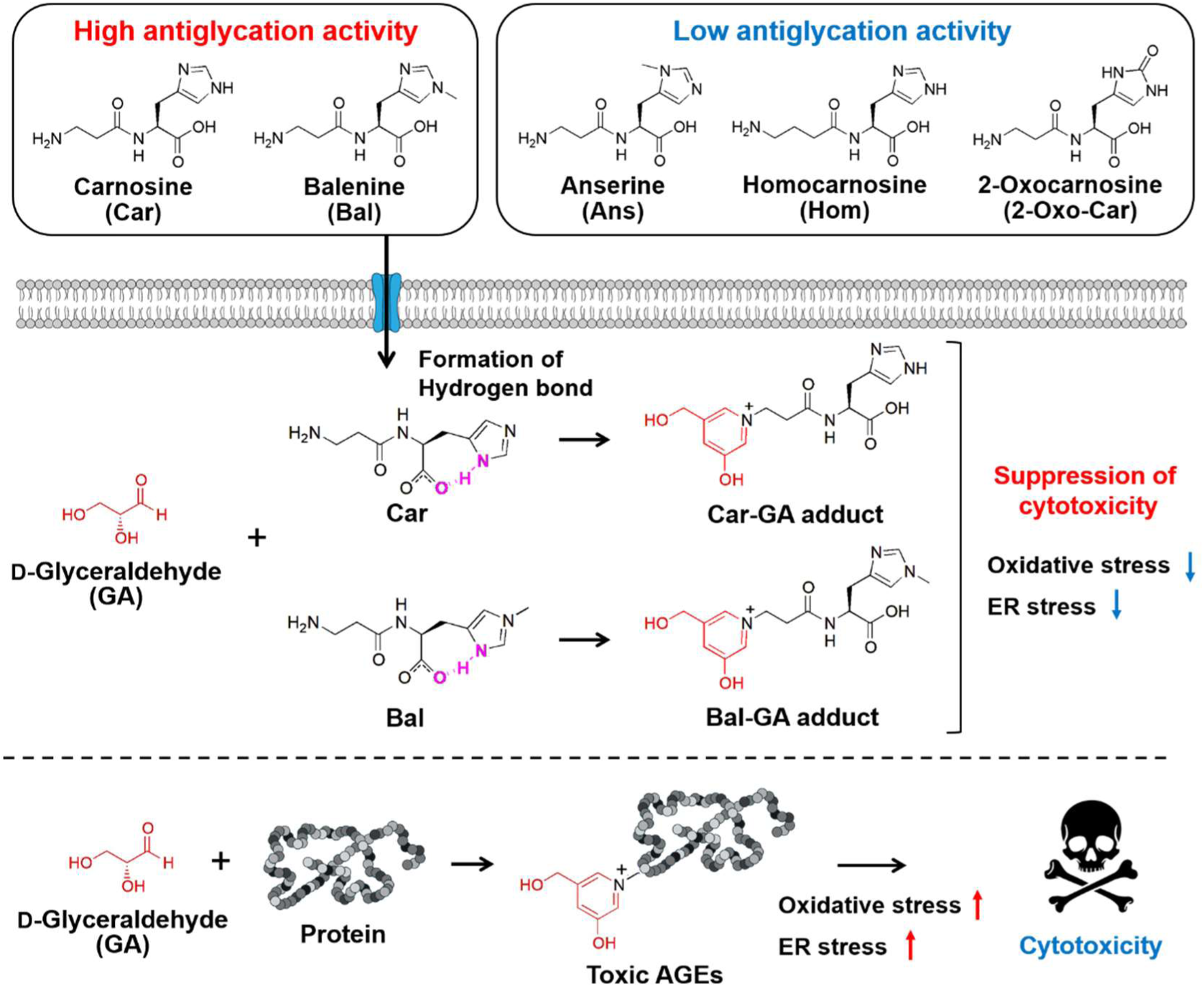
Schematic illustration of the GA-induced protein glycation and neuronal cell death suppression mechanisms of Car and Bal.

## Conclusions

We elucidated the molecular mechanism through which Car and Bal specifically scavenge GA, a precursor of TAGE, via their terminal amino groups. *In silico* analysis suggests that the selective capture process is regulated by intramolecular hydrogen bonding of IDPs. Importantly, as confirmed by western blot and qPCR analyses, the formation of GA-IDP adducts significantly attenuates intracellular AGE accumulation and cytotoxicity, and suppresses cellular responses associated with oxidative and ER stress. These results provide insight into the molecular mechanisms of neuroprotection against GA by IDPs and a rational strategy for the design of next-generation neuroprotective molecules with optimized chemical reactivity.

## Methods

### Materials

Car, Ans, Bal, Hom, 2-oxo-Car, His, *β*Ala, ammonium formate, acetonitrile (HPLC grade), 100 mM phosphate buffer (HPLC grade), Dulbecco’s Modified Eagle’s Medium (D-MEM) Ham’s F-12, bovine serum albumin (BSA), LDH cytotoxicity assay kit, Sepasol®-RNA I Super G, and Hanks’ balanced salt solution (HBSS) (+), and 1x phosphate-buffered saline (PBS) were obtained from Nacalai Tesque, Inc. (Kyoto, Japan). GA was purchased from Sigma Aldrich Japan (Kyoto, Japan). Ac-Car was purchased from Cayman Chemical Company (Ann Arbor, MI, USA). Anti-GA-pyridine antibody was purchased from Cosmo Bio Co. Ltd. (Kyoto, Japan). Anti-*β*-actin antibody was purchased from Medical and Biological Laboratories Co. Ltd. (Tokyo, Japan).

### HPLC and LC–MS measurements for Hom reagent

HPLC was performed using a SHIMADZU HPLC system (Shimadzu Corporation, Kyoto, Japan). IDP reagents were separated using a COSMOSIL 3PBr (3.0 mm I.D. x 250 mm, particle size; 3 µm, Nacalai Tesque) column^53, 54^ with 100 mM phosphate buffer (pH 7.0) using an isocratic mode at a flow rate of 1.0 mL/min at 30°C. UV detection was done at 220 nm and 250 nm. LC–MS analysis was conducted using a Nexera Lite HPLC system (Shimadzu) coupled with a single quadrupole mass spectrometer (LCMS–2050, Shimadzu). Hom reagent was analysed on a COSMOSIL 3PBr column (3.0 mm I.D. x 150 mm, particle size: 3 µm, Nacalai Tesque) using 20 mM ammonium formate under isocratic mode, at a flow rate of 0.3 mL/min During the LC–MS measurements, the nebulizing gas flow rate was set to 2.0 L/min, the drying gas flow rate to 5.0 L/min, the heating gas flow rate to 7.0 L/min, and the desolvation temperature was maintained at 450 ◦C. IDPs were detected at *m*/*z* 227.2 (Car), *m*/*z* 241.2 (Ans, Bal, and Hom), and *m*/*z* 257.1 (2-oxo-Hom). Data acquisition was performed in the positive ion mode with a detector voltage of 1.0 kV.

### Cell culture

SH-SY5Y cells (ECACC, accession numbers: 94030304, UK) were cultured in D-MEM Ham’s F-12 supplemented with 15% fetal bovine serum (FBS, Corning Inc., NY, USA). The cells were cultured in 6 cm dishes (Corning Inc.) at 37°C in a 5% CO_2_ atmosphere.

### Cell viability assay

SH-SY5Y cells (6.2 x 10^4^ cells) were seeded into 24-well plates and cultured at 37°C in a 5% CO_2_ atmosphere for 48 h. The culture medium was removed, and the cells were washed with HBSS (+). Next, the cells were treated with a mixture of 450 µL of D-MEM Ham’s F-12 containing 2% FBS, 50 µL of IDPs or 2-oxo-Car (20 mM) in 1x PBS, and 4 μL of GA (100 mM) at 37°C in a 5% CO_2_ atmosphere for 48 h. After incubation, 50 μL of Cell Count Reagent SF (Nacalai Tesque) was added to each well and incubated at 37°C in a 5% CO_2_ atmosphere for 2 h. The absorbance was measured at 450 nm (Abs_450 nm_) derived from formazan using a UV spectrophotometer (Infinite 200 PRO M Plex, Tecan, Männedorf, Switzerland). The cell viability was calculated as follows: (Abs_450 nm_ of the cells treated with GA and IDPs or 2-oxo-Car – Abs_450 nm_ of blank sample)/(Abs_450 nm_ of the cells with IDPs and 2-oxo-Car – Abs_450 nm_ of blank sample) x 100%.

### LDH assay

SH-SY5Y cells (6.2 x 10^4^ cells) were seeded into 24-well plates and cultured at 37°C in a 5% CO_2_ atmosphere for 48 h. The culture medium was removed, and the cells were washed with HBSS (+). Next, the cells were treated with a mixture of 450 µL of D-MEM Ham’s F-12 containing 2% FBS, 50 µL of IDPs or 2-oxo-Car (20 mM) in 1x PBS, and 4 μL of GA (100 mM) at 37°C in a 5% CO_2_ atmosphere for 48 h. After incubation, 100 μL of culture supernatant and fresh medium were added to a 24-well plate. Then, 100 μL of the substrate solution was added to each well and incubated in the dark at room temperature for 20 min. After incubation, 50 μL of stop solution was added to each well, and the absorbance was measured at 490 nm using a UV spectrophotometer (Infinite 200 PRO M Plex).

### Confirmation of the binding ability of IDPs or 2-oxo-Car to GA

For the binding assay, 20 µL of 100 mM phosphate buffer (pH 7.4), 20 µL of 100 mM GA, and 20 µL of 20 mM IDPs or 2-oxo-Car were mixed with 140 µL of ultrapure water. After a 24-h incubation at 37°C, the samples were analysed by HPLC, which was performed using a SHIMADZU HPLC system (Shimadzu). IDPs and IDP-GA adducts were separated on a COSMOSIL 3PBr column (3.0 mm I.D. × 150 mm, particle size: 3 µm, Nacalai Tesque) using 20 mM ammonium formate as solvent A in an isocratic mode, at a flow rate of 0.4 mL/min and a column temperature of 40°C. Between samples, the column was washed with 20 mM ammonium formate/acetonitrile (8/2, v/v) for 5 min, followed by equilibration with mobile phase (20 mM ammonium formate) for 15 min. UV detection was done at 210 nm, and the injection volume was 0.5 µL. The reaction rates of each IDP and 2-oxo-Car to GA were calculated as follows: reaction rates (%) = 100−[(peak area of the samples with GA/peak area of the samples without GA) x 100].

### Characterization of IDP-GA adducts by LC–MS measurements

Generated IDP-GA adducts were purified by HPLC. IDP-GA adducts were separated and purified on a COSMOSIL 3PBr column (4.6 mm I.D. × 250 mm, particle size: 3 µm, Nacalai Tesque) using 20 mM ammonium formate as solvent A under isocratic mode, at a flow rate of 1.0 mL/min and a column temperature of 40°C. The purified IDP-GA adducts were allowed to evaporate using a Smart Evaporator C1 (BioChromato, Inc., Kanagawa, Japan). The residue was then dissolved in ultrapure water. LC–MS analysis was conducted using a Nexera Lite HPLC system coupled with LCMS–2050 (Shimadzu). The purified IDP-GA adducts were analysed on a COSMOSIL 3PBr column (3.0 mm I.D. × 150 mm, particle size: 3 µm) using 20 mM ammonium formate as solvent A under isocratic mode, at a flow rate of 0.4 mL/min and a column temperature of 40°C. Between each sample analysis, the column was washed with a solution of 20 mM ammonium formate/acetonitrile (8/2, v/v) for 5 min, followed by equilibration with the mobile phase (20 mM ammonium formate) for 15 min. During the LC–MS measurements, the nebulizing gas flow rate was set to 2.0 L/min, the drying gas flow rate to 5.0 L/min, the heating gas flow rate to 7.0 L/min, and the desolvation temperature was maintained at 450°C. Data acquisition was performed in the positive ion mode with a detector voltage of 1.0 kV. Fragmentation patterns of the IDPs and IDP-GA adducts were recorded in positive ion mode across three collision energies (CEs; 100, 140, and 150 V), with the most distinct mass spectra obtained at the optimal CE for each compound.

### Evaluation of the antiglycation activity of IDPs and 2-oxo-Car on GA-induced glycation of BSA

BSA was dissolved in 1x PBS to a concentration of 5 mg/mL. Next, 10 µL of BSA solution was placed into a 1.5 mL centrifuge tube and mixed with 60 µL of ultrapure water, 10 µL of 10x PBS, 10 µL of 5 mM GA, and 10 µL of 20 mM IDPs or 2-oxo-Car or amino acids (His and *β*Ala). The reaction mixtures were incubated for 24 h at 37°C. Next, 30 μL of purified water, 50 µL of sample buffer solution containing 2-ME (2x) for SDS-PAGE (Nacalai Tesque), and 20 µL of the sample were mixed and heated at 95°C for 5 min. Glycation of BSA induced by GA and the inhibitory effect of the IDPs and 2-oxo-Car against glycation were assessed by SDS-PAGE followed by CBB staining and western blotting using anti-GA-pyridine antibody.

### Evaluation of the antiglycation activity of IDPs and 2-oxo-Car against GA-induced intracellular protein glycation

SH-SY5Y cells (2.8 x 10^5^) were seeded into 6-well plates and cultured at 37°C in a 5% CO_2_ atmosphere for 48 h. The culture medium was removed, and the cells were treated with a mixture of 1.8 mL of D-MEM Ham’s F-12 containing 2% FBS, 200 μL of 20 mM IDPs or 2-oxo-Car or amino acids (His and *β*Ala), and 16 μL of 100 mM GA at 37°C in a 5% CO_2_ atmosphere for 24 h. The cells were washed three times with 1 mL of PBS. Next, 500 μL of cell lysis buffer (Nacalai Tesque) was added to the culture plate, and the plate was shaken slowly for 5 min. The lysate and pellet were scraped using a cell scraper and transferred to a 1.5 mL microtube, which was placed on ice for 15 min and centrifuged at 10,000 x g at 4°C. Finally, the supernatant was mixed with sample buffer solution containing 2-ME (2x) for SDS-PAGE and heated at 95°C for 5 min. The protein concentration of each sample was adjusted to 200 μg/mL using a BCA assay kit (Nacalai Tesque) before mixing with sample buffer. Glycation of intracellular proteins induced by GA and the inhibitory effect of IDPs and 2-oxo-Car against glycation were assessed by SDS-PAGE followed by CBB staining and western blot analysis using an anti-GA-pyridine antibody.

### CBB staining

BSA solution or cell lysates were subjected to SDS-PAGE (10% Bullet PAGE Plus Precast Gel, Nacalai Tesque) and the gel was washed three times with purified water for 5 min. Next, the gel was shaken in CBB Stain One (Nacalai Tesque) for 1 hour at room temperature. The gel was then washed with purified water for 16 h and the bands were detected using a ChemiDoc Touch MP Imaging System (Bio-Rad Laboratories, Hercules, CA, USA).

### Western blot analysis

BSA solution or cell lysates were subjected to SDS-PAGE and transferred to polyvinylidene difluoride (PVDF) membranes (Merck Millipore, Nottingham, UK) using Bullet Semi-dry Transfer One (Nacalai Tesque) and the Trans Blot Turbo Transfer System (Bio-Rad Laboratories). Non-specific binding to the membrane was blocked using Bullet Blocking One (Nacalai Tesque) for 5 min. The PVDF membranes were washed and incubated for 16 h at 4°C with anti-GA-pyridine (Cosmo Bio Co., 1:1,000) or anti-*β*-actin (Medical and Biological Laboratories Co., Tokyo, Japan, 1:1,000) antibodies in 0.1% t-Tris-buffered saline (t-TBS) (Nacalai Tesque). *β*-Actin was used as an internal control. After washing, the membranes were incubated with horseradish peroxidase-conjugated mouse IgG (Nacalai Tesque, 1:10,000) in 0.1% t-TBS. The immunoreactive bands were detected using the ChemiDoc Touch MP Imaging system (Bio-Rad Laboratories) and Chemi-Lumi One Super (Nacalai Tesque). The intensity of the bands was quantified using Image Lab software (BioRad Laboratories).

### qPCR analysis

Total RNA was isolated from cells using Sepasol®-RNA I Super G based on the manufacturer’s protocol. cDNA was synthesized using the PrimeScript^TM^ RT Master Mix (Perfect Real Time) (Takara Bio Inc., Shiga, Japan). qPCR was performed using a Dice Real Time System III thermal cycler (Takara Bio) and THUNDERBIRD® Next SYBR® qPCR Mix (TOYOBO Co. Ltd., Osaka, Japan). Samples were quantified using the *ΔΔ*Ct method and normalized to *ATCB* (actin beta) levels to quantify relative mRNA expression. The qPCR primer sequences are listed in Table S1.

### In silico analysis

To determine the most stable conformation of each IDP and 2-oxo-Car, a conformational search study was performed using MOE software (Chemical Computing Group Inc., Canada). The coordinates of the IDPs and 2-oxo-Car were generated using ChemDraw and Chem3D software (PerkinElmer Inc., Waltham, MA, USA). The conformational search calculation was performed using the LowMode MD method with the MMFF94x force field.

### Statistical analysis

All data are expressed as the mean ± standard deviation (SD). Statistical difference among multiple groups was calculated with one-way ANOVA, followed by Tukey’s post hoc test. Groups sharing the same letter are not significantly different, whereas those with different letters are significantly different. *p*-values < 0.05 were considered statistically significant.

## Author contributions

Yasunari Yamada: conceptualization, investigation, methodology, validation, data curation, formal analysis, visualization, writing—original draft preparation, writing—review and editing. Koji Hashida: methodology, writing—original draft preparation, writing—review and editing. Kohei Hayashi: methodology, writing—review and editing. Kenji Yoshimochi: supervision, writing—review and editing. Tsunehisa Hirose: supervision, writing—review and editing. Motoshi Shimotsuma: funding acquisition, supervision, writing—review and editing. Yoshio Hamada: formal analysis, writing—original draft preparation, writing—review and editing. Kenji Usui: formal analysis, supervision, writing—review and editing. Naoki Yokoyama: resources, writing—review and editing. Toshiaki Hara: resources, writing—review and editing. Satoru Nishino: resources, writing—review and editing. Hideaki Kakeya: supervision, writing—review and editing. Shozo Tomonaga: supervision, writing—review and editing. Makoto Ozaki: conceptualization, investigation, methodology, validation, visualization, supervision, project administration, writing—original draft preparation, writing—review and editing.

## Data availability

The data supporting this article have been included as part of the supplementary information (SI).

## Supplementary data

**Fig. S1.**
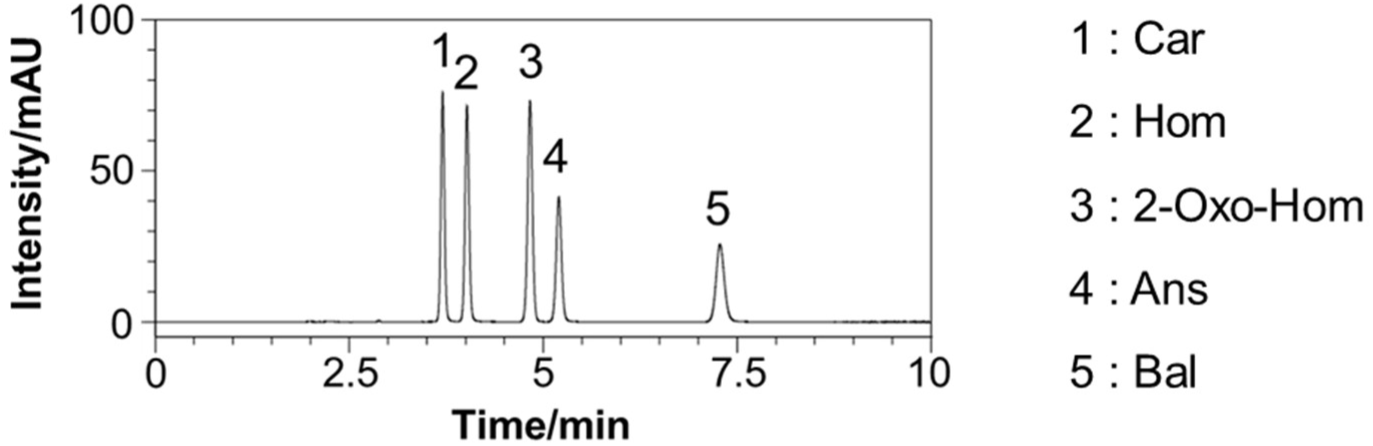
HPLC chromatogram of Car, Ans, Bal, Hom, and 2-oxo-Hom (0.2 mg/mL). The analysis was performed using a COSMOSIL 3PBr (4.6 mm I.D. × 250 mm, particle size: 3 µm). The mobile phase was 100 mM phosphate buffer (pH 7.0) in isocratic mode for 10 min at a flow rate of 1 mL/min and a column temperature of 30°C. The injection volume was 1 µL, and UV detection was performed at 220 nm.

**Fig. S2.**
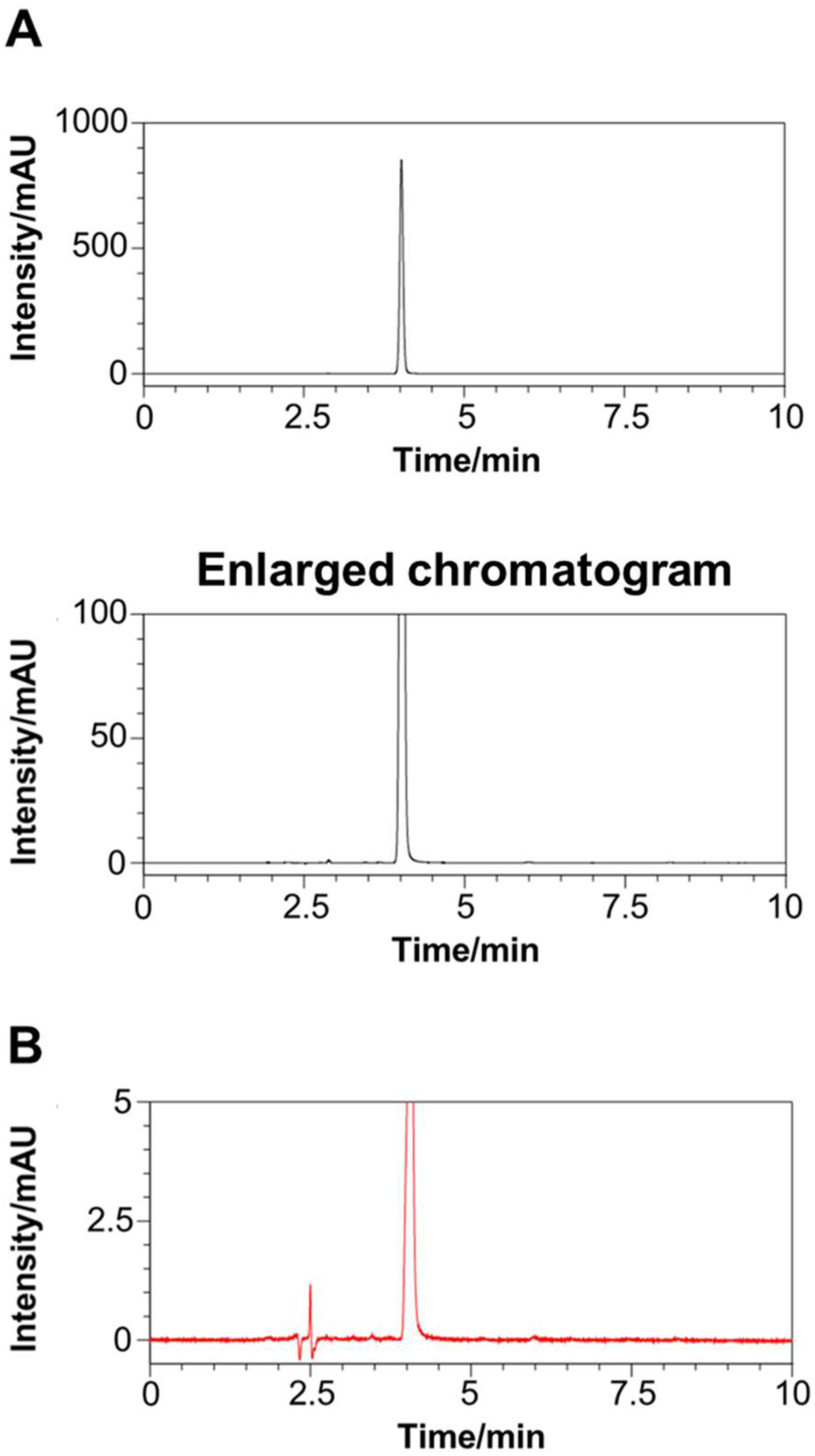
HPLC chromatograms of Hom (A: 1 mg/mL and B: 20 mg/mL). The analysis was performed using a COSMOSIL 3PBr (4.6 mm I.D. × 250 mm, particle size: 3 µm). The mobile phase was 100 mM phosphate buffer (pH 7.0) in isocratic mode for 10 min at a flow rate of 1 mL/min and a column temperature of 30°C. The injection volume was (A) 2.5 µL and (B) 5 µL, and UV detection was performed at (A) 220 nm and (B) 250 nm.

**Fig. S3.**
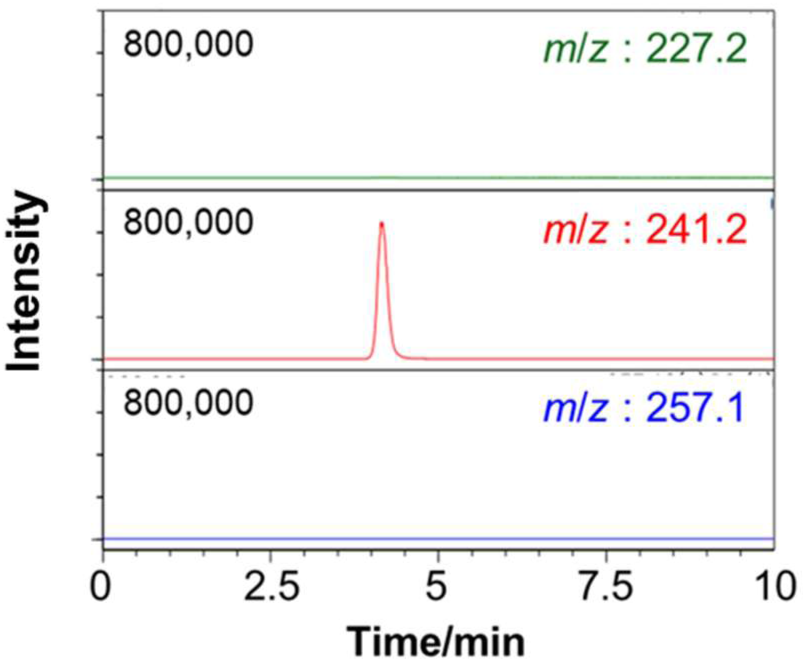
LC-MS chromatogram of Hom (10 µg/mL). The analysis was performed using a COSMOSIL 3PBr (3.0 mm I.D. × 150 mm, particle size: 3 µm). The mobile phase was 20 mM ammonium formate in isocratic mode for 10 min at a flow rate of 0.3 mL/min and a column temperature of 30°C. The injection volume was 0.5 µL.

**Fig. S4.**
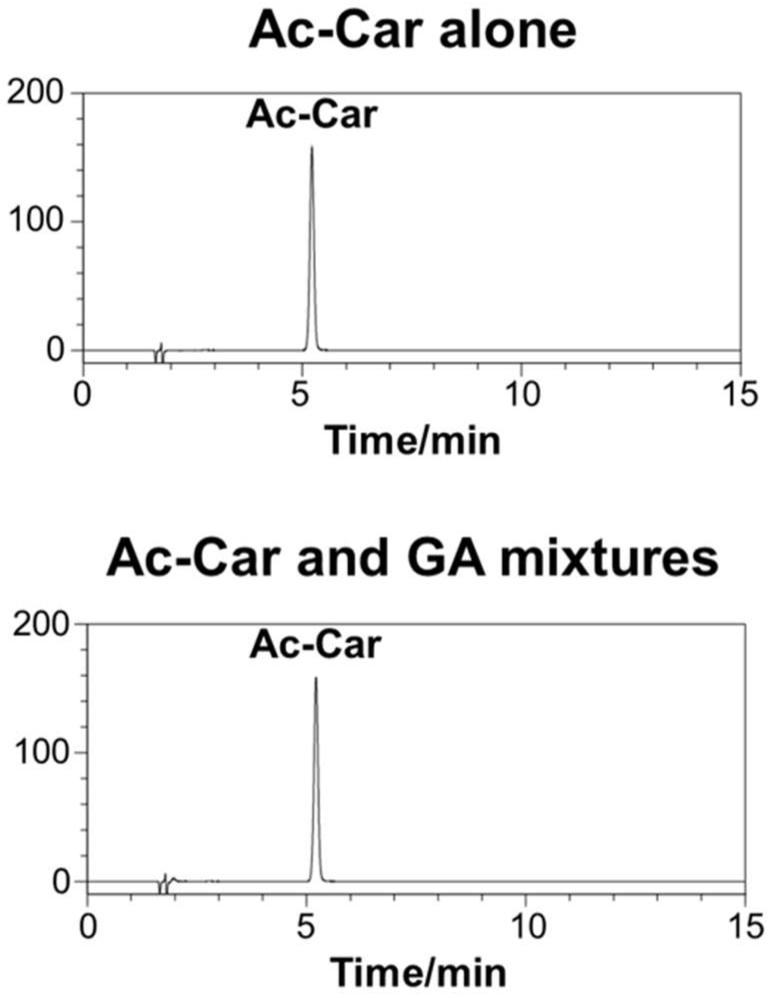
HPLC chromatograms of the samples following incubation with Ac-Car alone and Ac-Car with GA for 24 h at 37°C. The analysis was conducted using a COSMOSIL 3PBr (3.0 mm I.D. × 150 mm, particle size: 3 µm). The mobile phase was 20 mM ammonium formate in isocratic mode for 15 min at a flow rate of 0.4 mL/min and a column temperature of 40°C. The injection volume was 0.5 µL. UV detection was performed at 210 nm.

**Fig. S5.**
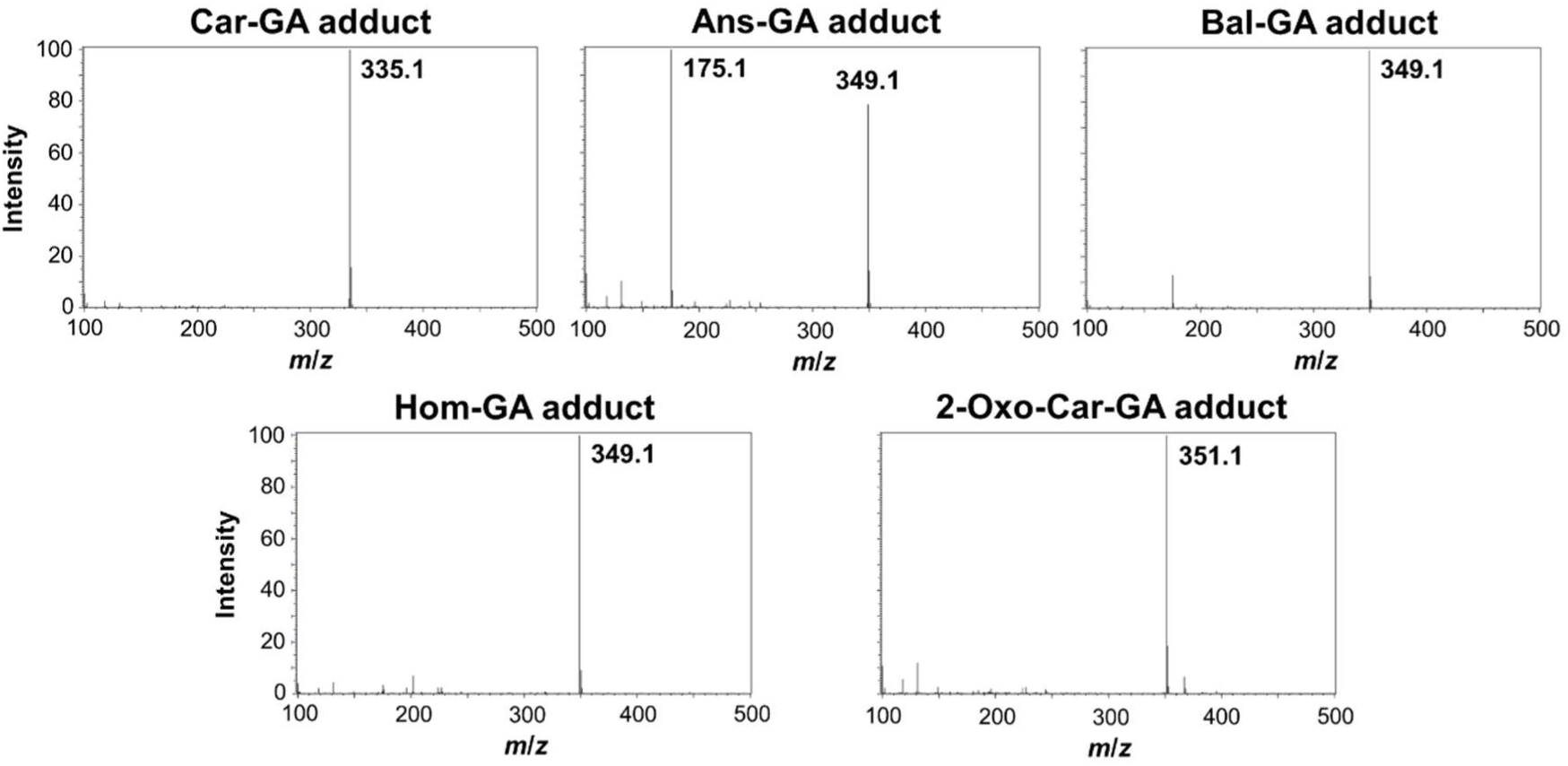
MS spectra of the IDP-GA and 2-oxo-Car-GA adducts.

**Fig. S6.**
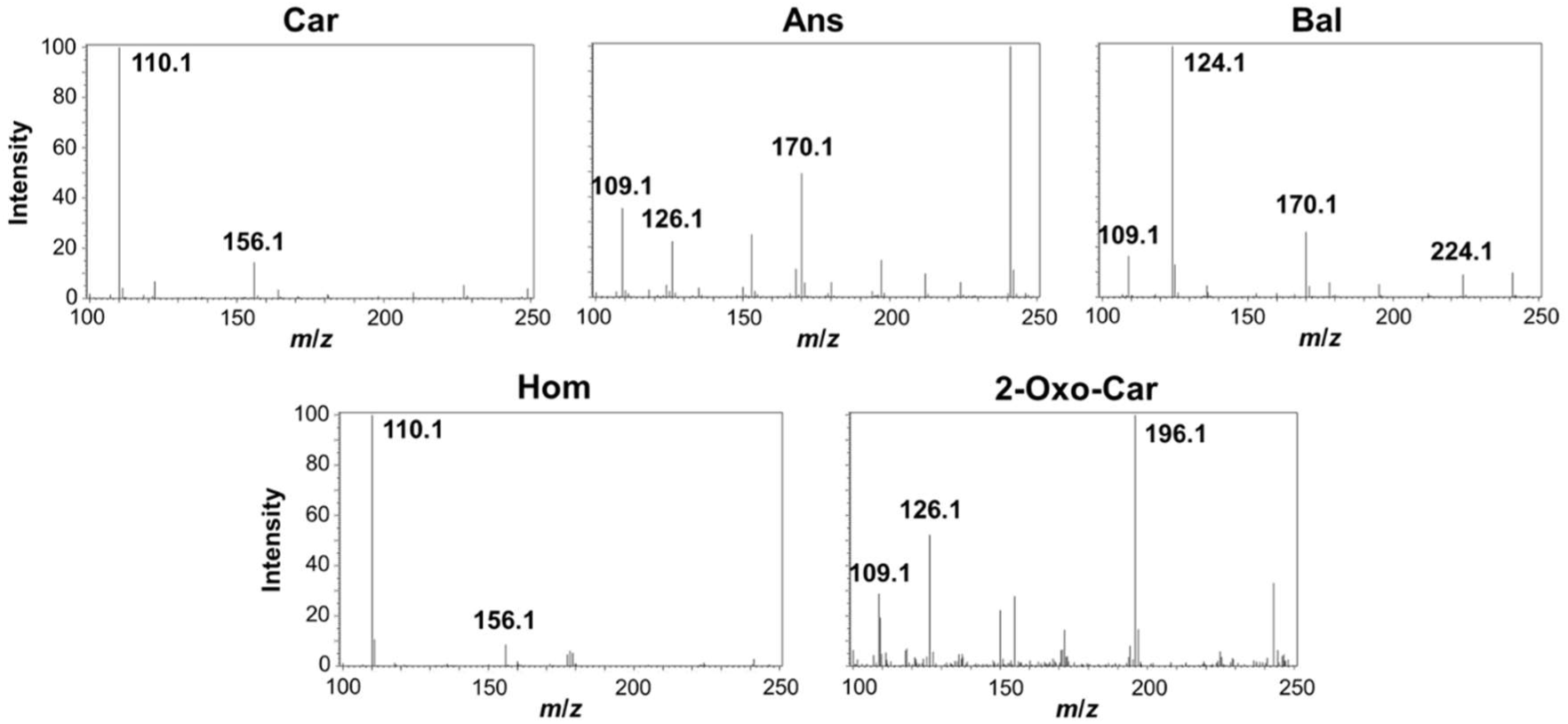
MS/MS spectra of Car, Ans, Bal, Hom, and 2-oxo-Car.

**Fig. S7.**
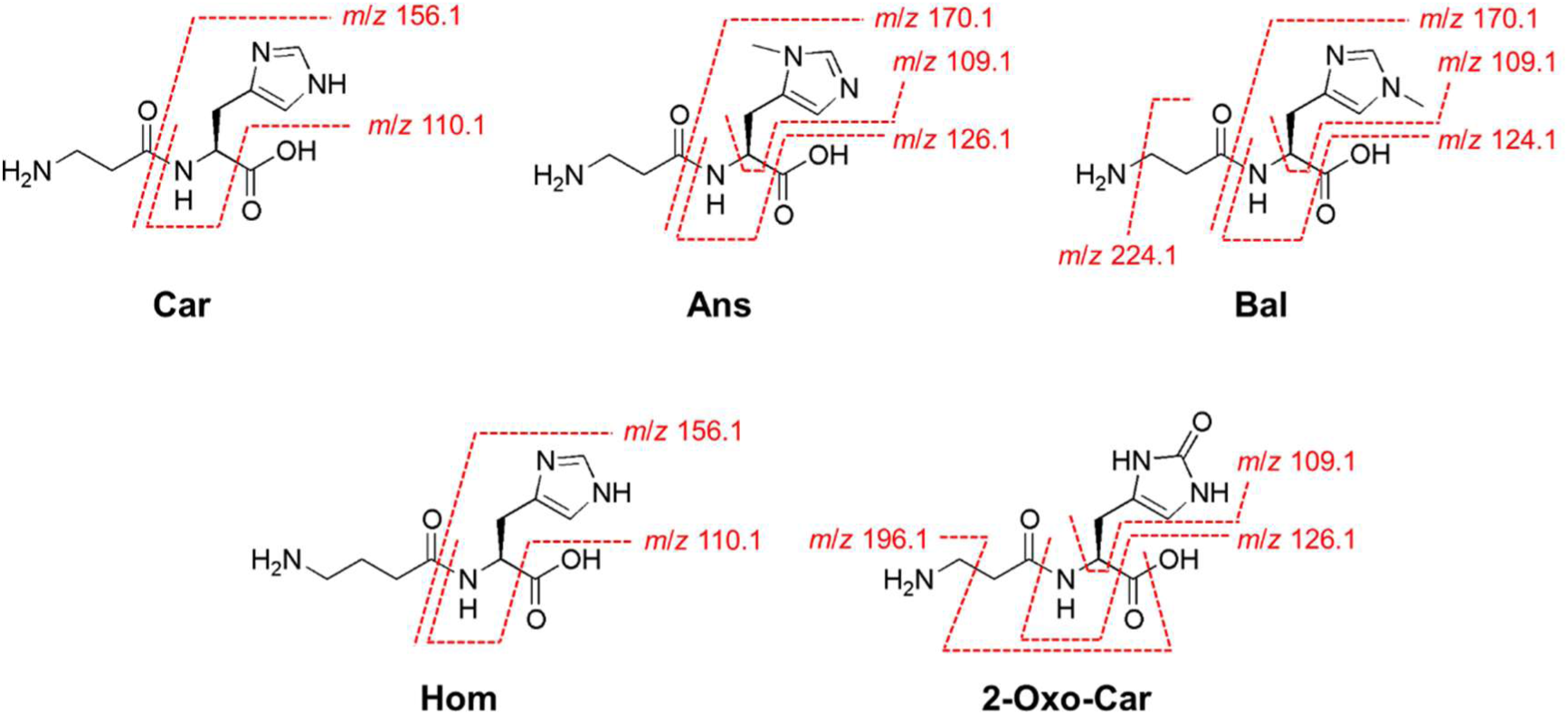
Chemical structures of the fragment ions of Car, Ans, Bal, Hom, and 2-oxo-Car. Cleavage sites are indicated by dashed lines.

**Fig. S8.**
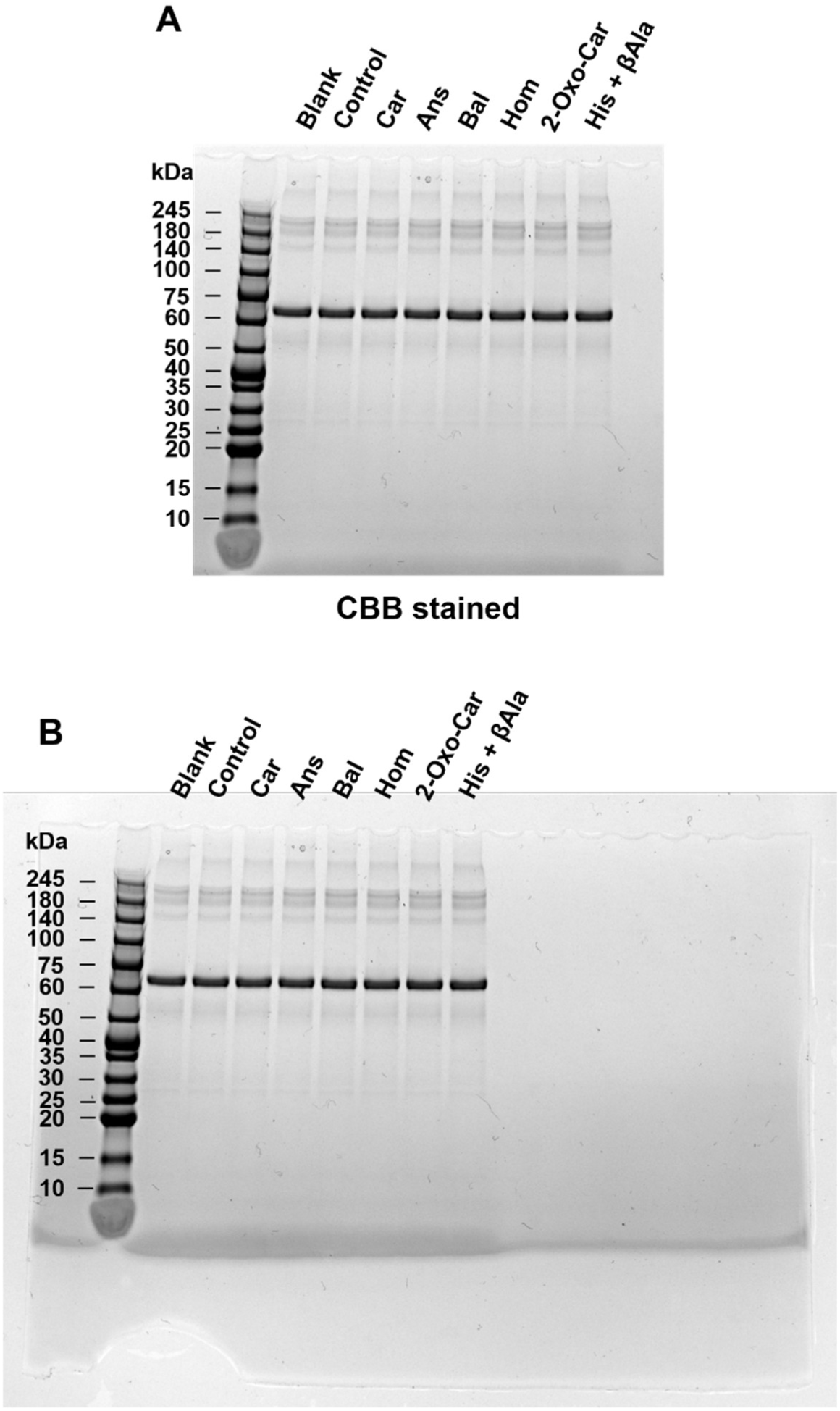
SDS-PAGE images (A: enlarged image, B: raw data) of BSA after incubation of GA with each IDP, 2-oxo-Car, or His and *β*Ala for 24 h at 37°C.

**Fig. S9.**
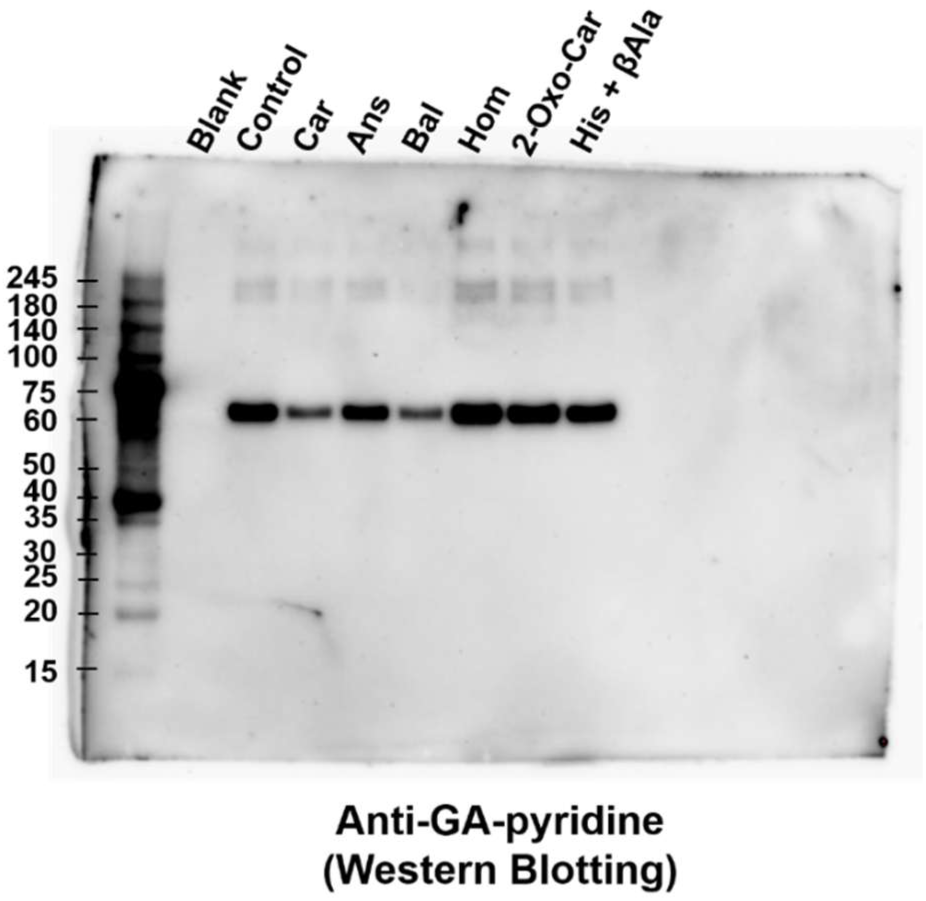
Western blot image (raw data) of BSA after incubation of GA with each IDP, 2-oxo-Car, or His and *β*Ala for 24 h at 37°C.

**Fig. S10.**
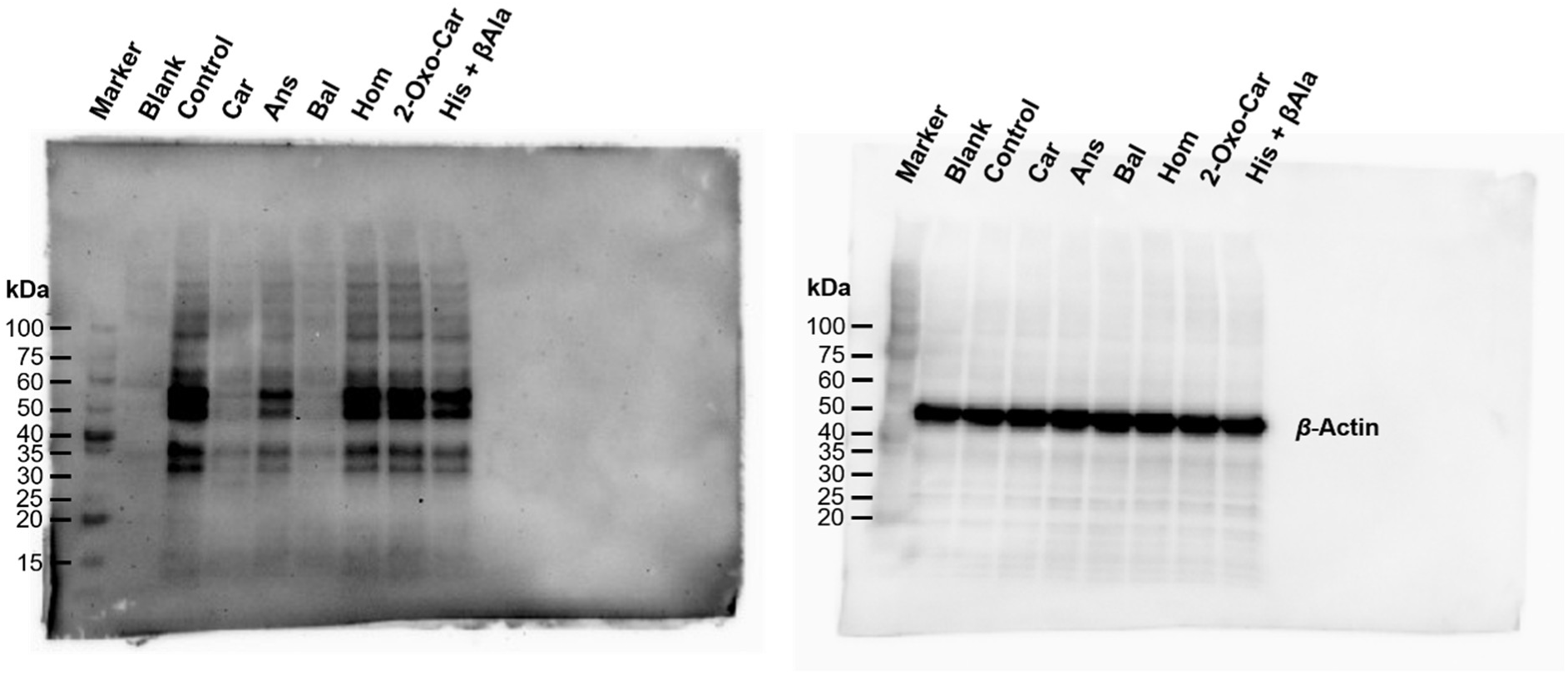
Western blot analysis of GA-induced intracellular protein glycation inhibition of IDPs, 2-oxo-Car, and His and *β*Ala.

**Fig. S11.**
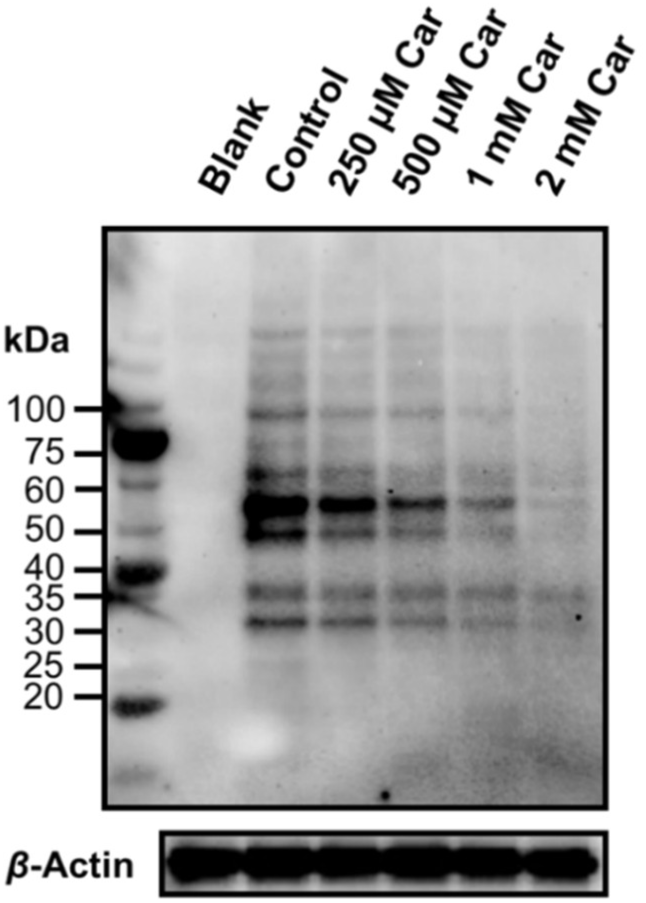
Western blot analysis of GA-induced intracellular protein glycation inhibition with increasing Car concentrations.

**Fig. S12.**
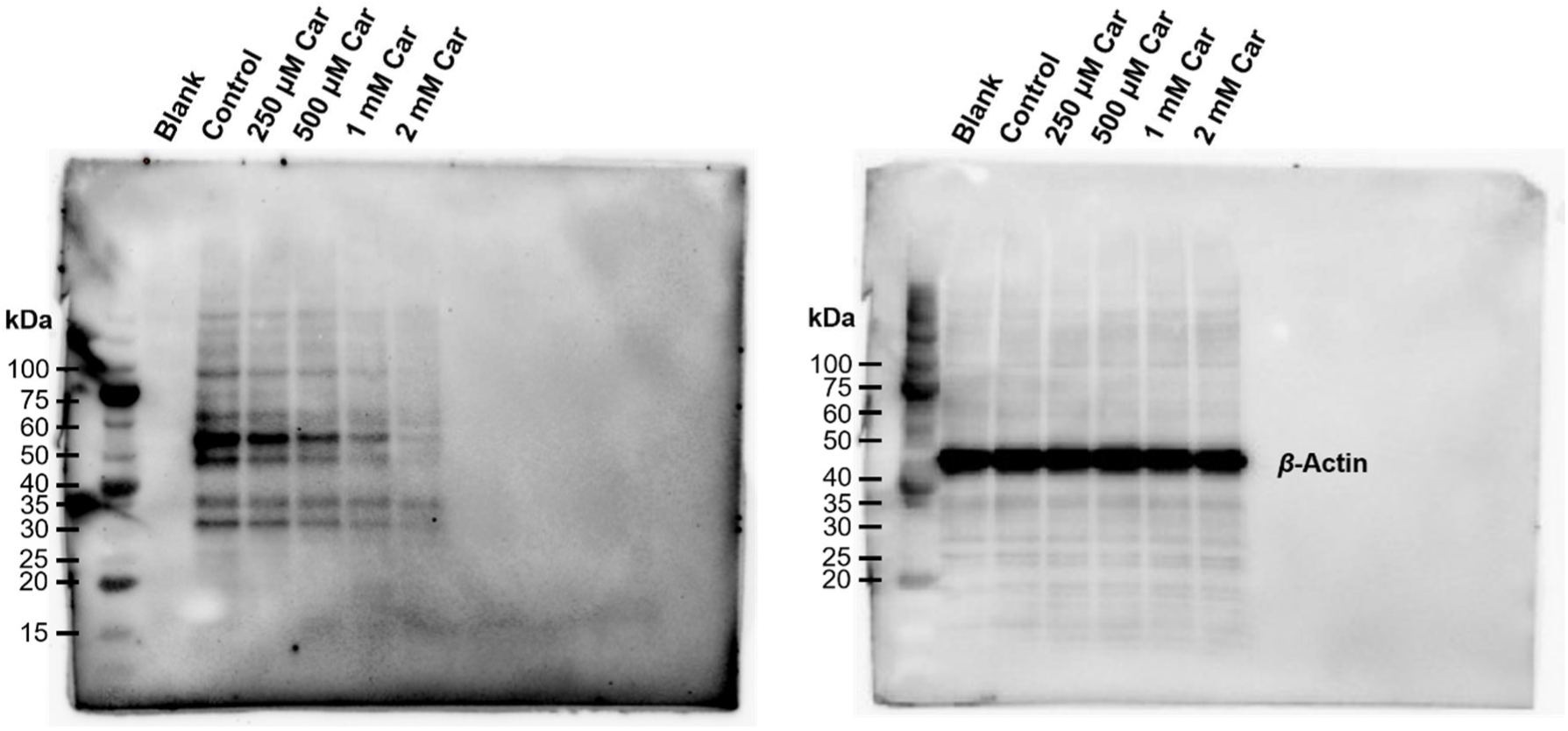
Western blot analysis (raw data) of GA-induced intracellular protein glycation inhibition at increasing Car concentrations.

**Fig. S13.**
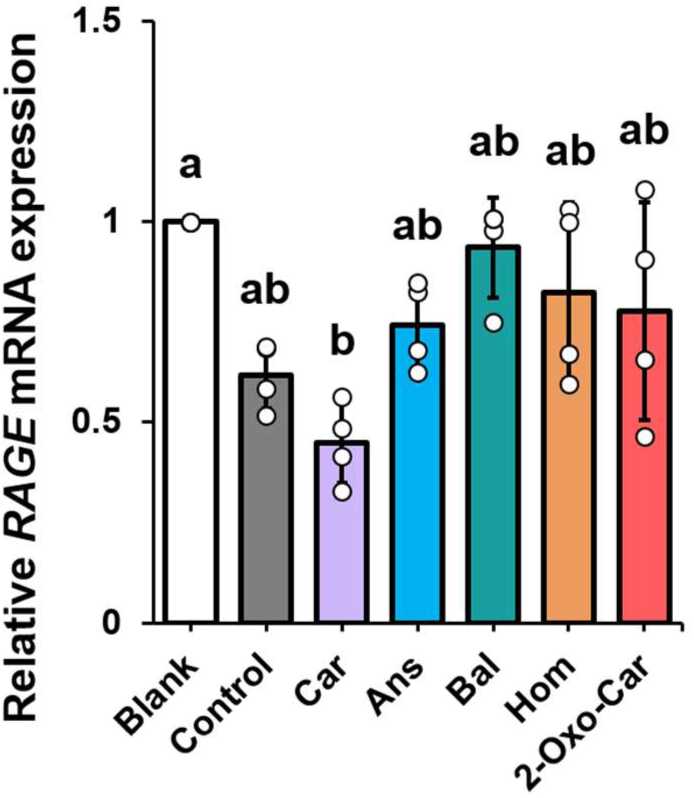
SH-SY5Y cells were treated with 800 μM GA and 2 mM IDPs or 2-oxo-Car for 24 h. *RAGE* expression was measured by qPCR. The data are presented as the mean ± SD (N = 4). Different letters indicate statistically significant differences among groups based on a one-way ANOVA followed by Tukey’s post hoc test (p < 0.05).

**Fig. S14.**
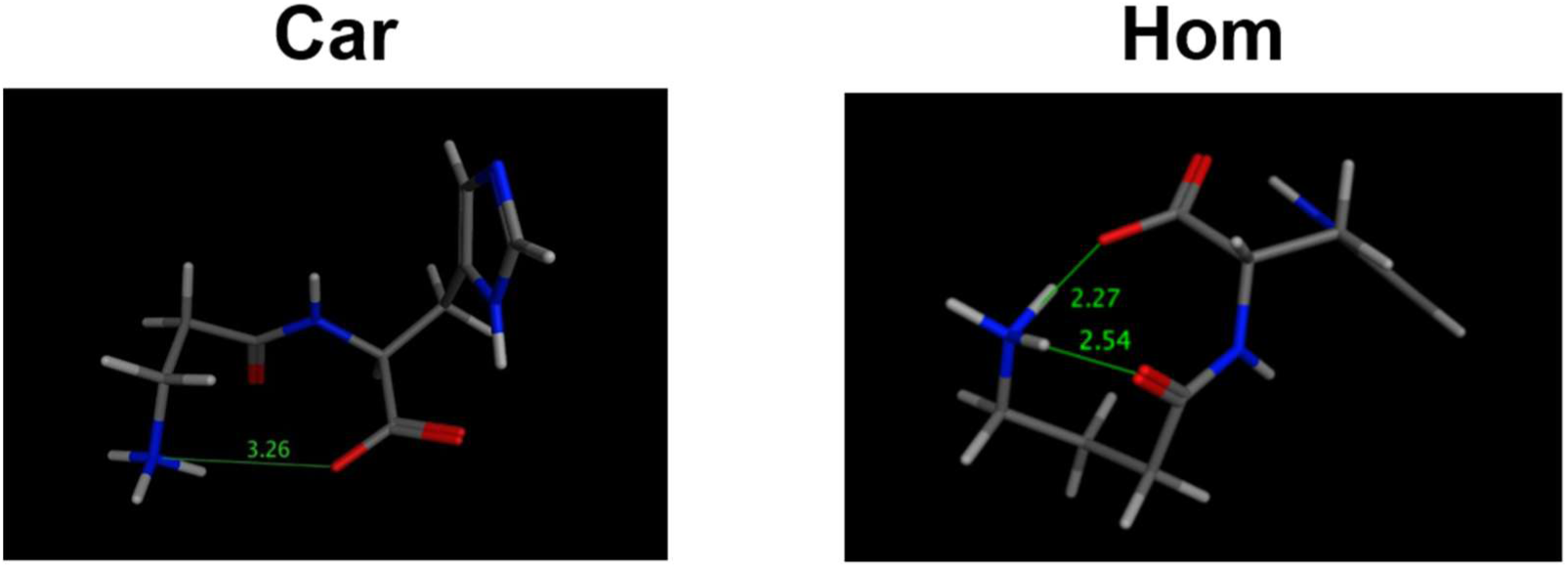
Interatomic distances of hydrogen bonds between C-terminal carboxyl group and N-terminal amino group of Car and Hom.

**Table S1.**
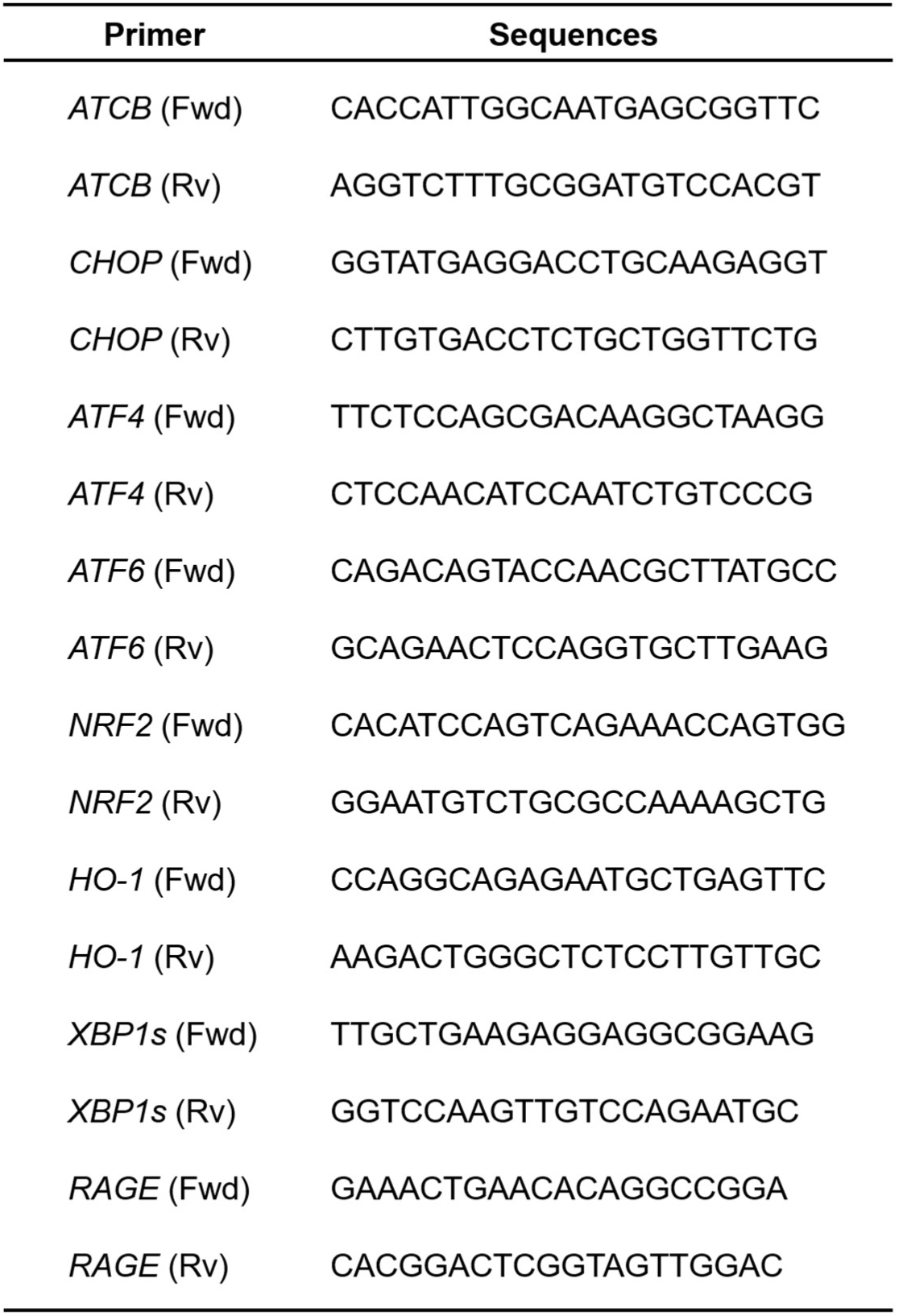
Primer sequences used for qPCR.

